# Deep Mechanistic Models reveal pathway-extrinsic drivers of mammary MAPK signalling heterogeneity

**DOI:** 10.64898/2026.07.30.741759

**Authors:** Giacomo Fabrini, Fabian Fröhlich

**Affiliations:** Dynamics of Living Systems Laboratory, The Francis Crick Institute, London, UK

## Abstract

Cells sense and respond to their environment through signalling pathways, and the dynamics of these pathways shape cell fate even within genetically identical populations. Two largely separate computational traditions describe this behaviour: mechanistic differential-equation models and representation-learning methods. Mechanistic models encode pathway topology and kinetics but cannot easily represent variation arising outside the modelled pathway. Representation learning, instead, maps genome-wide measurements onto low-dimensional manifolds but offers no mechanistic account of how the resulting cell states execute their functions. Reconciling these views, explaining signalling heterogeneity in a manner that is at once data-driven and mechanistically interpretable, has remained difficult. Here we introduce deep mechanistic models (DMMs), which couple semi-supervised representation learning to an ordinary-differential-equation model of EGFR/MAPK signalling, trained end-to-end so that the learnt representation and mechanistic parametrisation inform each other. Applying DMMs to multiplexed signalling data from 63 breast cancer cell lines, we show that the models generalise to held-out cell lines and attribute most heterogeneity to pathway-extrinsic factors, namely baseline ERBB2 activation and a Ca²⁺/p38 signalling axis, rather than to variation in core MAPK components. Where the models fail, the discrepancies pinpoint rare signalling-altering mutations and recurrent programmes, including a putative AMPK– BRAF MEK-inhibitor-resistance axis and a cytoskeletal programme. We further find that mechanistic integration of EGFR receptor levels reshapes the learnt representation, rendering a molecular and a systems-level account of the same data equivalent in predictive power. DMMs thus offer a general framework for fusing mechanism with learning, naturally extendable to further modalities such as imaging, and simultaneously turn model failure into a systematic route to discover and evaluate candidate biology.

## Introduction

Signalling enables cells to sense and respond to their environment, with the dynamics of information processing shaping cell fate. A canonical example is the EFGR/MAPK pathway (Lavoie et al., 2020), which is activated when extracellular ligands such as Epidermal Growth Factor (EGF) bind receptor tyrosine kinases (RTKs) such as EGFR (ERBB1) or ERBB2 (HER2/neu). Ligand binding induces receptor dimerisation and autophosphorylation, recruitment of adaptor proteins (e.g. GRB2/SOS), and activation of RAS. RAS then initiates a conserved three-tiered kinase cascade, RAF, MEK, and ERK. Activated ERK phosphorylates a wide range of substrates both in the cytoplasm and nucleus, including transcription factors and downstream kinases such as p90^rsk^ (ribosomal S6 kinase), which relay signals to regulate gene expression, protein synthesis, and cell growth. The pathway encodes information not only in signal amplitude but also in temporal features such as pulse frequency and duration (Avraham and Yarden, 2011; Ender et al., 2022; Santos et al., 2007), which can differentially regulate proliferation, differentiation, or apoptosis.

Despite this well-defined pathway architecture, ERK signalling responses are highly heterogeneous across cells, even in isogenic cell populations (Kramer et al., 2022). Such heterogeneity, in amplitude and spatio-temporal dynamics, is essential for fate specification and functional specialisation in multicellular organisms (Gagliardi and Pertz, 2024; Lavoie et al., 2020). Such specialisation is exemplified in the terminal duct–lobular unit (TDLU), the functional unit of breast tissue and the primary site of origin for most breast cancers. The latter include ductal carcinomas, which arise from milk-transporting ducts, and lobular carcinomas, which originate from the milk-secreting acini. The TDLU comprises a bi-layered epithelium consisting of an inner layer of luminal epithelial cells that are responsible for milk production and secretion, and an outer layer of basal, myoepithelial cells lining the basement membrane and providing structural support and contractile function.

In mice, EGFR is essential for TDLU development (Sebastian et al., 1998; Sternlicht et al., 2005), while loss of ERBB2 gives rise to delayed and defective ductal growth (Jackson-Fisher et al., 2004). Even though these murine studies suggest these RTKs are mostly activated in stroma, comparative analysis highlights pronounced differences in cell type specificity across species (Morato et al., 2023) and provides evidence for autocrine activation of EGFR in epithelial cells (Morato et al., 2021). In humans, evidence is largely restricted to cell lines, where EGFR and ERK signalling is required for proliferation (Accornero et al., 2012; Morato et al., 2021), and ERBB2 and EGFR overexpression mutually inhibit one another, and prime differentiation (Ingthorsson et al., 2016). This mutual exclusivity in RTK expression was recapitulated in cancer cell lines (Gambardella et al., 2022) and mirrors the divergence between the intrinsic molecular subtypes used for patient stratification (Perou et al., 2000; Sørlie et al., 2001): ERBB2 defines the HER2-enriched subtype, whereas EGFR is expressed in a substantial fraction of basal-like tumours (Nielsen et al., 2004). Notably, ERBB2 is widely recognised as an oncogenic driver (King et al., 1985) that, when overexpressed, spontaneously dimerises and activates downstream signalling in a ligand-independent manner (Harari and Yarden, 2000) and can be targeted by anti-HER2 therapies like trastuzumab (Slamon et al., 2001).

This positions EGFR and ERBB2 not only as central players in molecular pathways relevant to healthy tissue development and diseases such as cancer, but also as key determinants of distinct epithelial cell states. We will not attempt to provide a universal definition of what cell states are, or how they differ conceptually from related notions such as lineage, cell type, or cell identity, as this has been discussed elsewhere (Morris, 2019). Instead, we wish to highlight what is of biological interest and what is most pertinent to this study: a systems-level description of recurring differences in macromolecular composition that underpin the functional specialisation of cells.

This conceptual divide between molecular descriptions of signalling pathways and systems-level descriptions of cell states is mirrored in computational approaches. Signalling pathways are commonly modelled using differential equations that aim to capture pathway topology and kinetics (Klipp et al., 2005). Heterogeneity is incorporated either through variation in the expression of pathway components (Adlung et al., 2017; Fey et al., 2015; Fröhlich et al., 2018a; Lara et al., 2023) or by introducing statistical variability in kinetic parameters (Adlung et al., 2021; Almquist et al., 2015; Dharmarajan et al., 2019; Erdem et al., 2022; Fröhlich et al., 2018b; Karlsson et al., 2015; Wang et al., 2026). The former approach is limited to pathway-intrinsic influences, while the latter is not data-driven, limiting biological interpretability and the ability to make individual-level predictions or identify biomarkers. In contrast, cell states are typically inferred using representation learning methods that map high-dimensional, genome-wide measurements onto low-dimensional manifolds ranging from simple methods like principal component analysis (PCA) or non-negative matrix factorisation (Lee and Seung, 1999) through non-linear methods like UMAP (McInnes et al., 2020) or t-SNE (Maaten and Hinton, 2008) to deep learning approaches (Eraslan et al., 2019). These unsupervised approaches are informed by variance in marker expression rather than biological function, and do not directly yield mechanistic insight.

Here, we introduce deep mechanistic models (DMMs), which bridge these two paradigms by implementing semi-supervised representation learning coupled to mechanistic models of signalling. In DMMs, high-dimensional molecular data are encoded into a low-dimensional latent representation using an autoencoder setup, commonly used in unsupervised learning approaches. This representation parameterises an ODE model of a signalling pathway, which acts as a supervised regression head to predict signalling dynamics. The model is trained end-to-end against experimental data, yielding representations that are both predictive and mechanistically interpretable.

In contrast to neural ODEs (Chen et al., 2018) or Universal Differential Equations (Rackauckas et al., 2021), the differential equations in DMMs are fixed and do not operate in latent space, making them mechanistically interpretable. Compared to Physics Informed Neural Networks (Raissi et al., 2019), the differential equation is integrated numerically, making DMMs applicable to partially observed systems.

Conceptually similar model architectures include physical bottleneck neural networks (Schmitt et al., 2024), conditional universal differential equations (de Rooij et al., 2025) and MultiVeloVAE (Li et al., 2025), but such models have not been employed for representation learning in the context of signalling networks.

Applying DMMs to signalling data from 63 breast cancer cell lines, we show that a single model can be both data-driven and mechanistically interpretable: it attributes most heterogeneity to pathway-extrinsic factors, baseline ERBB2 activation and a Ca²⁺/p38 axis, that a fixed MAPK model alone could not capture. Model failures pinpoint rare signalling-altering mutations and recurrent programmes, including a putative AMPK–BRAF MEK-inhibitor-resistance axis.

## Results

### Deep Mechanistic Model of EGFR/MAPK signalling generalises to held-out cell lines

To bridge representation learning and mechanistic modelling in practice, we developed deep mechanistic models (DMMs) of EGFR and MAPK signalling. The DMM architecture (Fig. 1A) takes high-dimensional molecular measurements (e.g. transcriptomic or proteomic profiles) as input features (blue) and encodes them as low-dimensional latent embedding (pink) using a neural network (grey). This latent representation is then mapped to parameter deviations (red) that, together with parameter averages (yellow), define the parameters of a mechanistic ordinary differential equation (ODE) model of EGFR and MAPK signalling.

**Figure 1.**
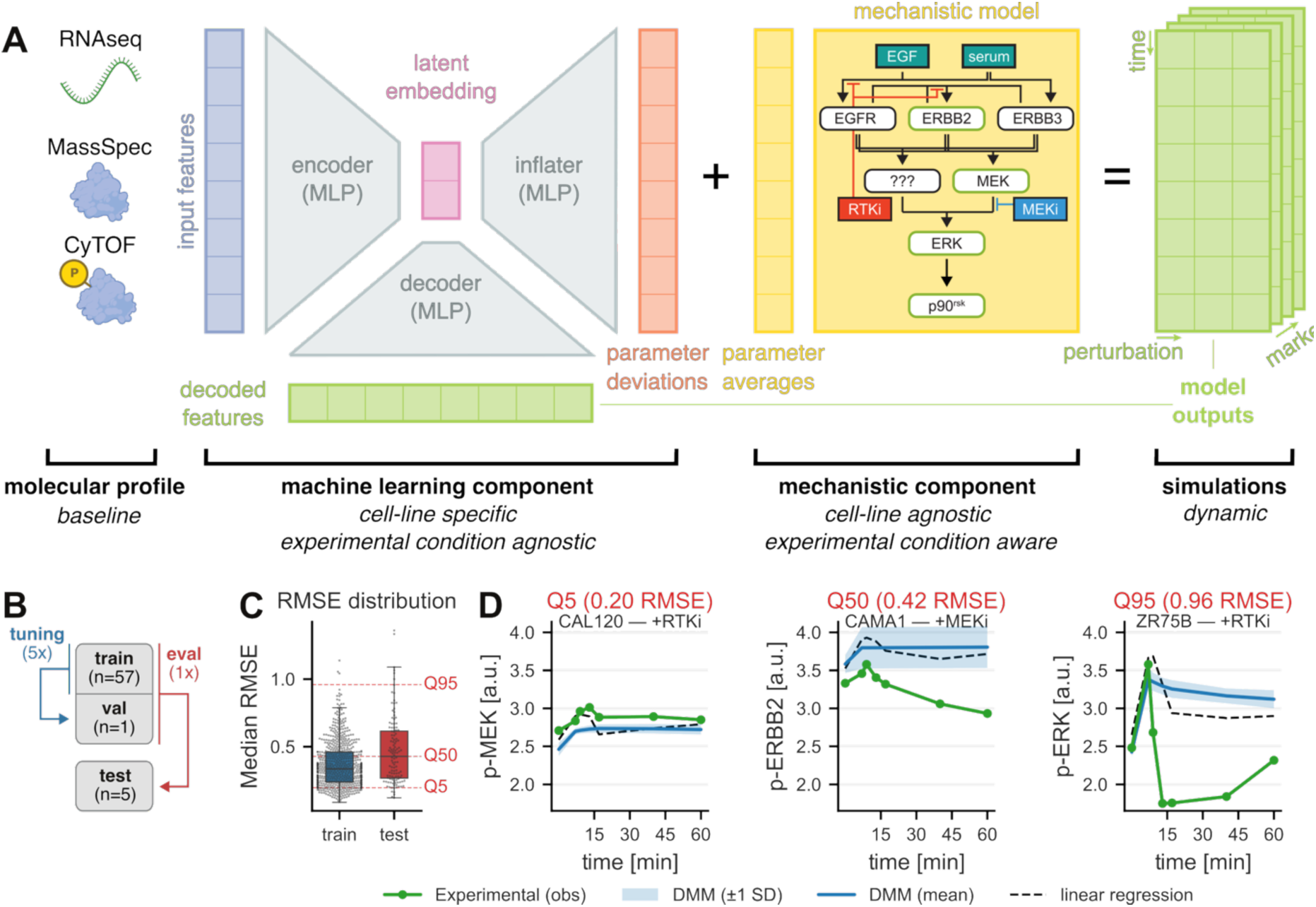
DMM architecture and representative model trajectories on the test set. (**A**) DMM model architecture including the machine learning and mechanistic components. **(B)** Training, validation and test split. Hyperparameter optimisation is performed in leave-one-out cross validation (LOOCV) over five worst-case validation lines in the training set. For evaluation on 5 held out test lines, the model is retrained on the full training set. **(C)** Distribution of per-combination (cell line × condition × observable, median over 10 training replicates) RMSE for the DMM on training (blue) and test (red) data. Boxplots show median and interquartile range; overlaid points represent individual (cell line × condition × observable) combinations. Dashed horizontal lines indicate the 5th, 50th, and 95th percentiles of the test RMSE distribution. **(D)** Representative predicted and observed time courses at the 5th (left), 50th (middle), and 95th (right) percentile of the test RMSE distribution. Circles indicate experimental measurements (mean across replicates); solid blue lines show DMM mean across training replicates; shaded band indicates ±1 SD across training replicates. Dashed black lines show linear regression baseline with elastic net regularisation. RMSE percentile and value are indicated above each panel.

The ODE model captures activation of the receptor tyrosine kinases (RTKs) EGFR, ERBB2, and ERBB3, together with downstream phosphorylation of MEK (MAP2K1, MAP2K2), ERK (MAPK1, MAPK3), and p90^rsk^ (RPS6KA1, RPS6KA2, RPS6KA3). It also accounts for MEK-independent (indicated by ???) ERK activation that has been suggested previously (Aksamitiene et al., 2010; Tognetti et al., 2021). It incorporates the effects of two small-molecule perturbations: the EGFR/ERBB2 inhibitor lapatinib (RTKi, red) and the MEK inhibitor PD184352 (MEKi, blue); and simulates signalling dynamics across time and perturbation conditions for a subset of phosphorylated markers in considered CyTOF panel (p-ERBB2, Tyr1196; p-MEK, Ser221; p-ERK Thr202/Tyr204; p-p90^rsk^ Ser380). These simulated dynamics constitute the model’s supervised output. In parallel the decoder reconstructs baseline features (unperturbed and at initial timepoint, green) from the same latent embedding.

The machine learning component was implemented in JAX (Bradbury et al., 2018), while the mechanistic model was implemented in PySB (Lopez et al., 2013) and AMICI (Fröhlich et al., 2021). To facilitate end-to-end differentiable training, we implemented custom vector–Jacobian products in pyPESTO (Schälte et al., 2023), thus enabling gradient-based training and ensuring that the learned latent representation captures information about dynamic, perturbation-dependent signalling responses.

We applied DMMs to CyTOF data from a panel of 63 breast cancer cell lines (Gabor et al., 2021; Tognetti et al., 2021), comprising four stimulation conditions: serum alone (single time point), EGF and serum stimulation either alone (10 time points, EGF condition) or with either the MEK inhibitor PD184352 (MEKi, +MEKi condition) or the EGFR and ERBB2 inhibitor lapatinib RTKi (+RTKi condition), where inhibitors were added 15 minutes prior to stimulation (7 time points). The dataset was split into a training set (58 cell lines) and a test set (5 cell lines) using the same partition as the 2019 Breast Cancer Signaling Challenge (Gabor et al., 2021). Extensive hyperparameter tuning was performed on the training set using leave-one-out cross-validation (LOOCV) over five worst-case validation lines (Fig. 1B). The five held-out cell lines in the training set were selected based on their deviation from the average response.

To assess generalisation performance, we trained a model with input features constructed from CyTOF readouts and evaluated its performance on training (blue, Fig. 1C) and test sets (red). We observed good generalisation performance, as indicated by similar root mean squared error (RMSE) values on the training set (0.34, blue) and the test set (0.43, red). Evaluation of predictions on the held-out cell lines (Fig. 1D) further confirmed strong agreement between model predictions (grey) and experimental measurements (green). Overall, these results demonstrate that DMMs can generalise effectively to previously unseen cell lines.

### DMMs achieve the lowest prediction error on multimodal data and can distinguish luminal and basal subtypes

To assess the information content of different molecular feature sets, we compared DMMs trained on transcript (RNA-seq; orange), protein (mass spectrometry, MassSpec; green), phosphoprotein (mass cytometry, CyTOF; blue), multimodal inputs combining transcript, protein and phosphoprotein levels (multimodal, red) and embeddings of the Multi-Omic Synthetic Augmentation foundation model (Cai et al., 2024) (MOSA; purple) (Fig. 2A, left). When predicting responses in unseen cell lines, all modalities except MOSA embeddings outperformed baseline mechanistic models trained on averaged data (negative control, dashed line), but none matched mechanistic models fitted to individual cell lines (positive control, dotted line), which define the upper bound on achievable DMM performance. Prediction error decreased with increasing regulatory proximity of the features to the signalling layer, although this trend may partly reflect technical biases. DMMs trained on multi-modal features performed best and clearly separated luminal (purple, Fig. 2B left) from basal (teal) cell lines in their latent embedding.

**Figure 2.**
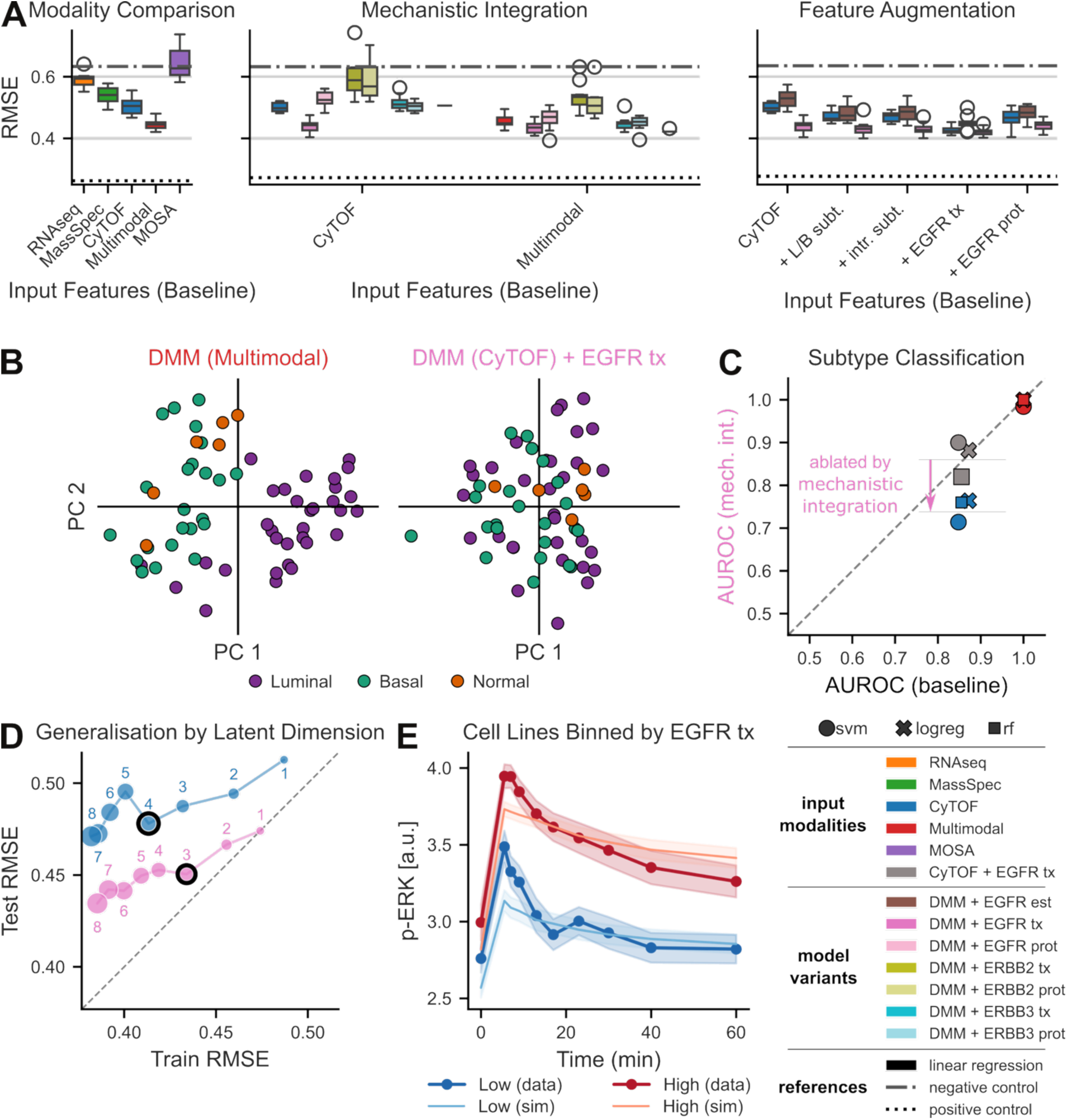
Deep Mechanistic Model performance across data modalities, RTK integrations, and feature augmentations. (**A, left)** Test RMSE of the base Deep Mechanistic Model (DMM) trained on five input data modalities: bulk RNA-seq (RNAseq, orange), mass spectrometry proteomics (MassSpec, green), mass cytometry (CyTOF, blue), multimodal (CyTOF combined with proteomics and transcriptomics, red), and the Multi-Omic Synthetic Augmentation (MOSA, purple) foundation model embeddings. Each bar represents the distribution across training replicates. **(middle)** Comparison of DMM variants incorporating mechanistic RTK (receptor tyrosine kinase) integrations for EGFR (pink), ERBB2 (green), ERBB3 (cyan) with transcript abundance (tx, dark shading) and protein abundance (prot, light shading) against the base DMM (blue) and an elastic-net linear regression baseline (black stripe), evaluated on held-out test data. Model variants are grouped by input data modality. **(right)** Effect of input feature augmentation on DMM test-set RMSE. The base DMM (CyTOF, blue), DMM with EGFR synthesis rate as free parameter (DMM + EGFR est, brown), and DMM with mechanistically integrated EGFR transcript levels (DMM EGFR tx, pink) are compared. Feature augmentation includes 1-hot encoded luminal/basal subtype (L/B subt.), intrinsic subtype (intr. subt.), EGFR transcript levels (EGFR tx) and EGFR protein levels (EGFR prot). Annotations based on (Marcotte et al., 2016) **(B)** Principal component analysis of DMM latent embeddings coloured by luminal (purple), basal (teal) or normal (orange) subtype for 58 breast cancer cell lines (training set, 30 luminal, 23 basal, 5 normal). Left: base DMM trained on multimodal input features; right: DMM with mechanistically integrated EGFR transcript levels (tx) trained on CyTOF input feature. Embeddings are aggregated across training replicates using PCA. **(C)** Comparison of luminal/basal subtype classification accuracy (AUROC) with (y-axis) and without (x-axis) mechanistic integration of EGFR transcript levels. Each point represents one classifier (support vector machine (svm): circle; logistic regression (logreg): cross; random forest (rf): square) evaluated with a given input feature set (CyTOF: blue, CyTOF+EGFR transcript levels: grey, multimodal: red). The dashed diagonal marks equal performance. **(D)** Train (x-axis) and test (y-axis) RMSEs for DMMs with different numbers of hidden units for the base DMM (blue) and DMM with mechanistic integration (pink). Black outline highlights the local minimum in test RMSE (base: 4; mechanistic integration: 3). **(E)** EGF-stimulated p-ERK dynamics stratified by EGFR transcript level into Low (blue) and High (red). Cell lines are split into EGFR-expression bins by median quantile. Solid lines with shading show the mean ± SEM across cell lines in each bin measured by CyTOF. Lighter-coloured lines show the corresponding model simulations averaged over training replicates and cell lines in each bin.

### Mechanistic integration of EGFR levels reshapes the latent representation at equal predictive power

Luminal and basal cell lines differ in ERBB2 and EGFR expression, receptors that are not only systems level subtype markers but also known molecular drivers of signalling heterogeneity (see Introduction). We next exploited this mechanistic role, incorporating receptor-level information directly in DMMs. Specifically, we trained DMMs with modified mechanistic models (Fig. 2A, middle) in which synthesis rates of EGFR (pink), ERBB2 (green), or ERBB3 (cyan) were rescaled based on measured transcript (tx, dark shading) or protein (prot, light shading) levels, following the individualisation approach employed in (Fröhlich et al., 2018a). For multimodal and CyTOF-based inputs, mechanistic integration of RTK levels only improved test performance for EGFR transcript levels.

The performance differences between transcript and protein-based mechanistic integration were surprising, given that ERBB2 and EGFR have been reported to show some of the highest transcript-to-protein correlations in pan-cancer studies (Nusinow et al., 2020). Comparing transcript and protein levels (Fig. S6) and examining peptide-level measurements (Fig. S7) across cell lines revealed that a small number of peptides dominates the estimated protein levels in cell lines with few transcripts, suggesting that the lack of predictive power is likely due to data artefacts.

Analysing the learnt latent structure for DMMs with mechanistically integrated EGFR transcript levels (Fig. 2B, right), we did not observe separation between luminal and basal cell lines for CyTOF-based DMMs, suggesting that mechanistic integration ablated learnt subtype structure. We confirmed this by a drop in subtype discrimination (Fig. 2C) with support-vector machine (svm, circle), logistic regression (logreg, cross) and random forest classifiers (rf, square) for CyTOF-features (blue) but not multimodal features (red). We hypothesised that mechanistic integration of EGFR levels constrains the effective latent space of DMMs. Consistent with this, analysis of prediction RMSEs across bottleneck sizes (Fig. 2D) showed a local minimum in prediction error at three latent dimensions when EGFR levels were mechanistically integrated, compared with four when they were not. Together, these observations suggest that, by injecting the subtype-correlated EGFR signal through the pathway parameters rather than leaving it to be learnt, mechanistic integration supplies one latent dimension directly, thereby ablating subtype structure and lowering the effective dimensionality.

To compare mechanistic integration of EGFR levels with data-driven approaches and assess their information content relative to subtype labels, we performed feature set augmentation (Fig. 2A, right) using one-hot encoded luminal/basal and intrinsic subtypes, as well as EGFR transcript and protein levels. To avoid limitations from model expressivity, we extended the mechanistic model with a scaling parameter for EGFR synthesis (EGFR est., brown). While most augmentations yielded modest performance gains, both the base (blue) and extended (brown) models remained significantly worse than models with mechanistically integrated EGFR levels (pink), except for EGFR transcript augmentation. In contrast to mechanistic integration, feature augmentation with EGFR transcript levels did not affect discriminatory power of embeddings for subtypes (grey, Fig. 2C).

To assess the functional consequence of EGFR expression, we grouped cell lines into high (red) and low (blue) transcript-level expression and investigated p-ERK dynamics following EGF stimulation for each group (thick lines, Fig. 2E). The high-expression group showed greater ERK phosphorylation than the low-expression group, a difference recapitulated by DMM simulations with mechanistically integrated EGFR transcript levels (thin lines). While this is consistent with the reported role of EGFR in setting EGF sensitivity (Fröhlich et al., 2023; Gerosa et al., 2020), it does not exclude other factors contributing to subtype-specific sensitivity.

Our findings demonstrate that, in contrast to feature augmentation, mechanistic integration reshapes the learnt latent structure, enabling switching between a molecular lens, in which EGFR drives signalling phenotypes through its molecular role within the pathway, and a systems lens, in which it acts as a marker distinguishing luminal and basal subtypes. Because both models achieve similar predictive power, we cannot claim that either representation is superior, only that DMMs enable downstream analysis under either lens.

### DMMs suggest baseline ERBB2 activation and Ca^2+^/p38 signalling as two major axes contributing to signalling heterogeneity

The lack of luminal–basal structure in the latent embedding of DMM models with CyTOF-based inputs prompted us to investigate what structure the model captures instead, and how it can be interpreted biologically. Neither visual inspection nor data-driven analyses revealed meaningful correlations with alternative cell-type annotations, such as MSI status, site of origin, or metastatic status (data not shown). We therefore turned to the inferred parameter deviations of the CyTOF-based DMM model with mechanistic integration.

Sensitivity analysis identified 6 out of 44 parameters with individually significant contributions to the training-set RMSE (one-sided one-sample t-test on the increase in RMSE after fixing each parameter at its population average, i.e. setting its deviation to zero, across n = 10 training replicates; Benjamini–Hochberg FDR < 0.05, Fig. 3A). Out of these 6 parameters, two also had significant, positive contributions to test set RMSE, which we subsequently refer to as major **heterogeneity axes**: those governing ERK to p90^rsk^ (*P1*, teal) and ERBB2 autoactivation (*P2*, orange). These parameter axes are highly correlated with other parameters (Fig. 3C) and the first three PCA embeddings of dynamic trajectories (Fig. 3D) that explain about 68% of variance. This confirms that these axes explain a major fraction of the heterogeneity observed in the data. Moreover, it demonstrates that even though these axes are mechanistically orthogonal, they are statistically entangled and therefore difficult to infer with purely statistical methods.

**Figure 3.**
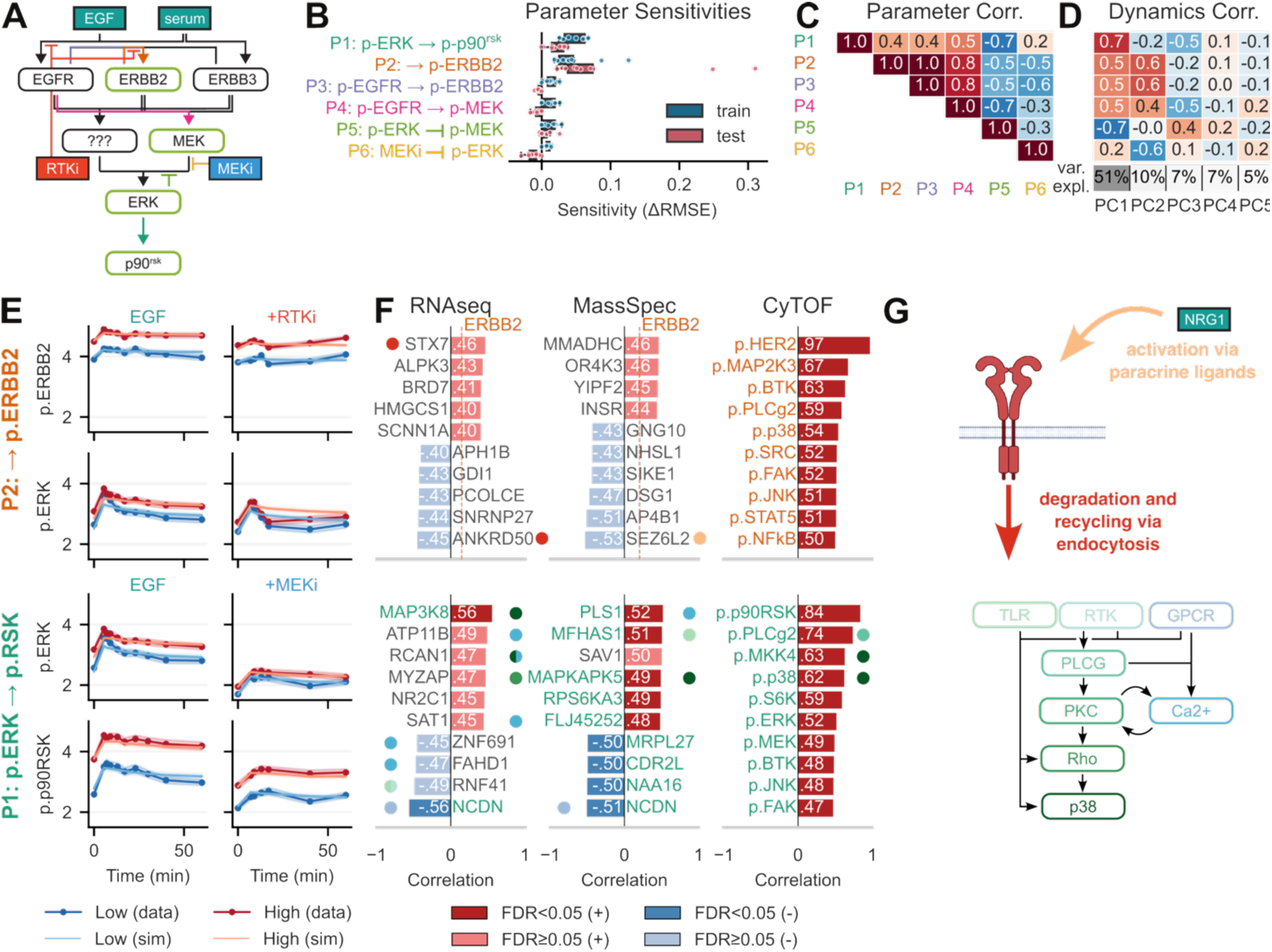
Mechanistically inferred EGFR–MAPK signalling parameters link to multi-omic cell-state identity. (**A**) Schematic illustration of model topology with arrows corresponding to sensitive parameters from (B) highlighted in the same colour. (**B)** Horizontal box-and-strip plots show the increase in root-mean-square error (ΔRMSE) upon individual ablation of the six most sensitive parameters (P1–P6) of the EGFR–MAPK mechanistic model, evaluated separately on training (dark blue) and held-out test (dark red) set. Parameters are ranked by mean sensitivity. Individual markers show variability across training replicates (n=10). **(C)** Upper-triangular Spearman rank correlation matrix between the six most sensitive parameters, computed across cell lines. Colour indicates positive (red) or negative (blue) correlations. **(D)** Spearman correlation between each inferred parameter and the principal components (PCs) of the full CyTOF dynamic data; variance explained per PC is shown below the heatmap. **(E)** Signalling dynamics stratified by inferred parameter. For each of two focal parameters: ERBB2 autophosphorylation rate (row 1; → p.ERBB2, shown in orange) and ERK→p90^rsk^ signal-transmission gain (row 2; p-ERK → p-p90^rsk^, shown in teal) cell lines are split into Low (blue) and High (red) groups at the population mean. CyTOF phosphoprotein abundance (mean ± SEM) is shown over time for EGF stimulation and a pathway inhibitor condition (+RTKi, top rows; +MEKi, bottom rows). Solid circles with shaded error bands: experimental data; solid lines with lighter shading: model simulations. **(F)** Multi-omic marker correlations. The ten markers most strongly correlated (Spearman ρ, Benjamini–Hochberg corrected) with each inferred parameter in RNA-seq transcriptomics (left), mass-spectrometry proteomics (middle), and CyTOF (right). Bars are colored by sign of correlation; opacity reflects significance (saturated = q < 0.05; pale = not significant). Dashed reference lines mark selected genes: ERBB2 (top row). Dots next to bars indicate association with biological process shown in (G). Employed model is CyTOF-based features with mechanistically integrated EGFR (tx) levels. Inferred parameters were z-score normalised and then averaged across training replicates (n=10). **(G)** Schematic illustration of cellular programmes identified in (F).

To validate these parameter axes, we grouped cell lines (Fig. 3E) into equally sized high (red) and low (blue) groups and compared group-averaged experimental data (thick lines with markers) with model simulations (thin lines). All parameters separate cell lines along either the source or target of the signalling step they describe, which is recapitulated by the simulations. Minor mismatches between data and simulations remain, such as transient pathway activation within the first 20 minutes under +RTKi conditions. However, as these discrepancies are consistent across groups, they do not affect interpretation of the parameter axes and likely reflect model limitations; lacking a clear mechanistic hypothesis of what could drive these dynamics, we did not pursue further model refinement. To biologically interpret these parameter axes, we correlated inferred parameter values with levels of transcriptomic, proteomic and CyTOF markers (Fig. 3F):

The *P2* axis captured a constant offset between parameter groups that persists under RTK inhibition (Fig. 3E, top *P2* panels). High baseline ERBB2 activation (red) was associated with more sustained ERK activation (Fig. 3E, bottom left *P2* panel), consistent with lower ERBB2 internalisation compared to EGFR (Baulida et al., 1996). Despite accounting for differences in internalisation, the model did not recapitulate this difference in signalling sustainment. The *P2* axis was strongly correlated with baseline p-ERBB2 levels (r=0.97, q=9.2×10^-29^ Benjamini–Hochberg adjusted p-value), indicating that phosphorylation dynamics are predominantly governed by basal activity. No statistically significant associations with transcriptomic or proteomic levels were observed; nevertheless, many correlations considerably exceeded those for ERBB2 transcript (r=0.15, vertical line) or protein (r=0.20) levels.

Among the top 10 correlated transcripts and proteins, only a few have mechanistically known links to ERBB2 signalling, where ANKRD50 (r=-0.45, q=0.9) and STX7 (r=0.46, q=0.9) participate in ERBB2 recycling and degradation. Interestingly, we observed a strong anti-correlation with SEZ6L2 (r=-0.53, q=0.76), a known BACE1 substrate. BACE1 is the sheddase that also processes the ERBB3/4 ligand NRG1 (Savonenko et al., 2008). Since the detected SEZ6L2 peptides derive from the shed and secreted ectodomain, a drop in inferred protein levels might indicate BACE1 shedding activity, which would be consistent with elevated ERBB2 phosphorylation via NRG1-driven heterodimer activation (Fig. 3G).

Conversely, the *P1* axis captured signal amplification between ERK and p90^rsk^, evidenced by comparable p-ERK levels across groups (Fig. 3E, top panels of *P1*), particularly under +MEKi conditions (Fig. 3E, top right *P1* panel), contrasted with a clear separation at the level of p-p90^rsk^ (bottom *P1* panels). We observed significant correlations with baseline p-p90^rsk^ (r=0.84, q=2.9×10^-16^) levels as well as p-PLCG2 (r=0.74, q=1.5×10^-9^), p-MKK4 (r=0.63, q=3.7×10^-7^) and p-p38 (r=0.62, q=4.5×10^-7^). At the transcriptomic level, we observed statistically significant correlations for TPL2/COT (MAP3K8, r=0.56, q=0.01) and Neurochondrin (NCDN, r=-0.56, q=0.01) as well as 13 proteins (PLS1, NHP2, MRPL3, BAG6, ANXA11, RPS6KA3, MAPKAPK5, FLJ45252, CDR2L, NAA16, NCDN, MFHAS1, MRPL27). Many of these genes participate in a Ca^2+^–PLCG–PKC–p38 signalling axis operating downstream of diverse transmembrane proteins including RTKs, TLRs and GPCRs (Lavoie et al., 2020; Lemmon and Schlessinger, 2010). Closer inspection revealed 10 genes among the top-10 most correlated transcripts and top-10 most correlated proteins (7 transcripts: MAP3K8, NCDN, ATP11B, RCAN1, MYZAP, FAHD1, RNF41; 4 proteins: PLS1, MFHAS1, MAPKAPK5, NCDN; see Table S4) that either contribute to this signalling axis or are involved in calcium binding. While this pathway has been extensively studied in neuronal, cardiac, and immune contexts, Ca²⁺-mediated signalling is also central to the function of mammary epithelial cells during lactation (Cross et al., 2014; Kelleher, 2022). Mechanistically, both non-canonical activation of p90^rsk^ by p38 (Zaru et al., 2015, 2007), and altered interactions with calcium-responsive scaffolds like KSR1 (Thines et al., 2024), IQGAP (Ren et al., 2008), or S100B (Hartman et al., 2014) could plausibly account for the observed differences in signal transduction. Collectively, these findings suggest the Ca^2+^/p38 signalling axis (Fig. 3G) may be a major regulator of signalling from ERK to p90^rsk^ in mammary cell lines.

Collectively, these findings suggest that MAPK signalling heterogeneity in mammary cell lines is governed by pathway-external factors, such as baseline ERBB2 activation and processing, and crosstalk with Ca²⁺/p38 signalling, rather than variation in core MAPK components. Notably, no strong correlations were observed with AKT-pathway markers (p-AKT^T308^, p-AKT^S473^), despite well-established MAPK-AKT crosstalk, although this may be the result of stimulation with saturating levels of EGF (Aksamitiene et al., 2012). More broadly, this demonstrates that DMMs complement the pathway-centric framing of mechanistic models with data-driven inference that reaches beyond the encoded topology, helping to discover candidate mechanisms for pathway-extrinsic regulation.

### Model failure identifies cell lines with rare signalling-altering mutations

Mechanistic models are considered most informative when they fail to reproduce experimental data, as such discrepancies can reveal gaps in biological knowledge. To evaluate the capacity of DMMs to uncover knowledge gaps, we analysed the distribution of training RMSE values across cell lines, observables, and conditions (Fig. 4A). RMSE values were generally lower for p-MEK and p-ERBB2 than for p-ERK and p-p90^rsk^, consistent with observations from purely statistical models (Gabor et al., 2021), likely reflecting the lower dynamic range for these markers. For p-ERK and p-p90^rsk^, error levels were particularly high under +RTKi conditions, representing a systematic failure to capture the transient activation of p-ERK following RTK inhibition, as previously discussed. We also observed elevated RMSE values under +MEKi conditions, with distinct error subgroups emerging across markers and cell lines: *error group E1* (yellow; HCC2185) exhibiting high error in p-ERBB2, p-ERK, and p-p90^rsk^; *error group E2* (green; DU4475) restricted to p-ERK; *error group E3* (purple; HCC1806, MCF12A, LY2, MDAMB157, 184B5, AU565, ZR751, HCC38, T47D, CAL148, HCC1187) affecting p-ERK and p-p90^rsk^; *error group E4* (magenta; MDAMB175VII, ZR7530) showing elevated error in p-p90^rsk^ and under +RTKi conditions.

**Figure 4.**
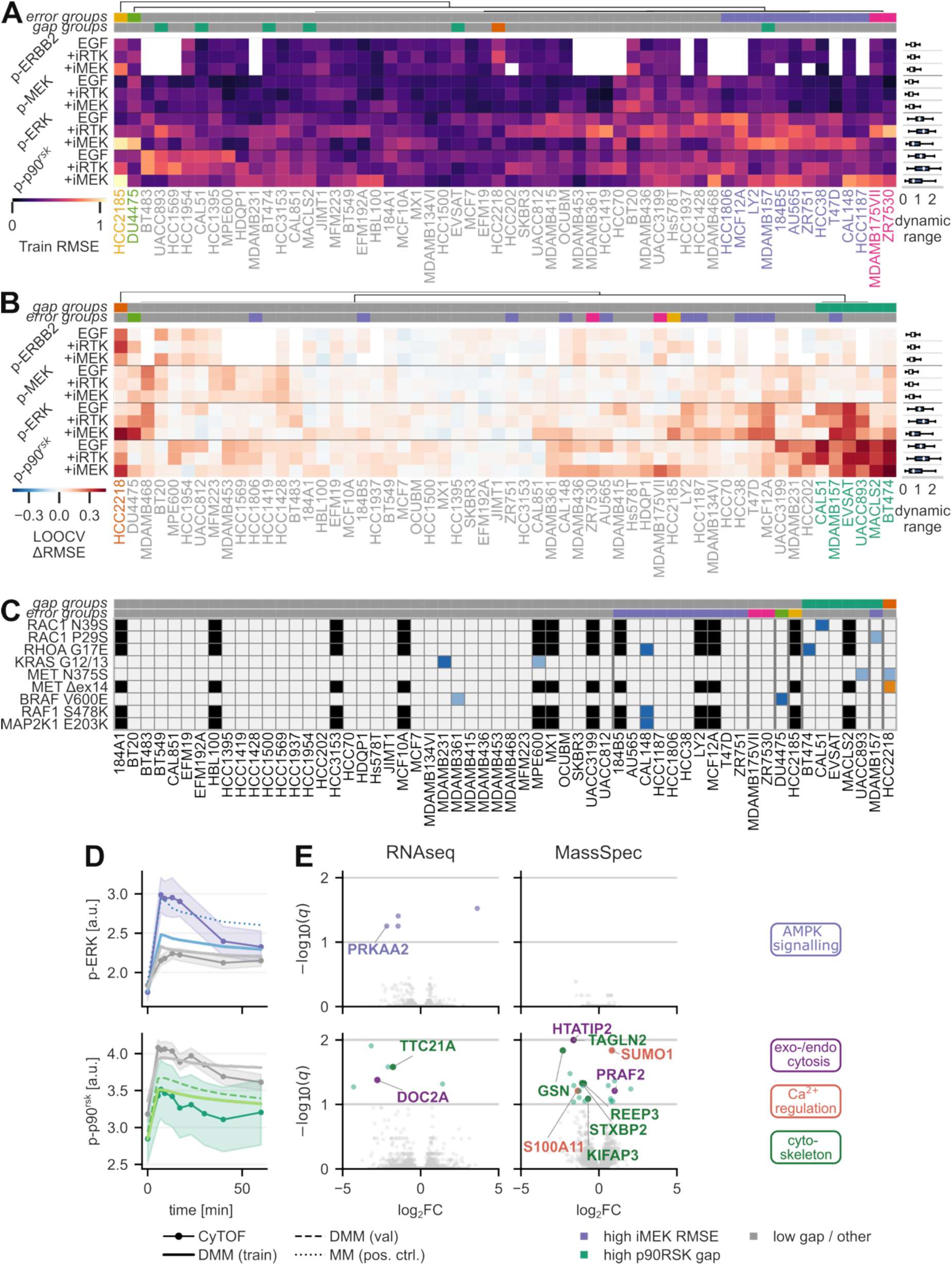
Cell line subgroup membership and omics associations with model generalisation gap. (**A**) Heatmap of training RMSE for p-ERBB2, p-MEK, p-ERK, and p-90^rsk^ under EGF stimulation, RTK inhibitor (+RTKi), and MEK inhibitor (+MEKi) conditions across breast cancer cell lines. Cell lines are ordered by hierarchical clustering (Ward linkage, dendrogram shown above). Coloured annotation strips below indicate ΔRMSE gap subgroup (top) and training RMSE subgroup (bottom) membership. Adjacent boxplot shows the distribution of dynamic range in data trajectories across cell lines. Colour bar indicates RMSE magnitude. White squares indicate missing data for p-ERBB2 measurements. **(B)** Heatmap of the generalisation gap (ΔRMSE = validation RMSE −training RMSE) from leave-one-out cross validation for the same observables, conditions, and cell line ordering as in (A). Colored annotation strips and adjacent boxplot as in (A). **(C)** Mutation status of selected variants in EGFR-MAPK pathway genes across cell lines ordered as in (A–B). Mutation data were retrieved from cBioPortal and the Cancer Models Project (CMP/COSMIC). Full-saturation colouring denotes variants confirmed by two or more independent sources; pale colouring denotes single-source variants; black squares indicate genes for which no mutation data were available. **(D)** Mean phosphoprotein trajectories under MEK inhibition for pERK (top) and under EGF stimulation for p90^rsk^ (bottom), stratified by subgroup membership. CyTOF measurements are shown as circles (mean ± s.e.m. across cell lines in each subgroup). Overlaid lines show DMM simulations from training cell lines (solid), DMM simulations for held-out cell lines from leave-one-out cross-validation (dashed), a mechanistic model positive control (dotted), and linear regression (with elastic net regularisation predictions for held-out cell lines (dash-dotted). **(E)** Volcano plots of differential expression between subgroup members and all other cell lines for the high MEKi RMSE subgroup (top row) and the high p90^rsk^ gap subgroup (bottom row), in transcriptomics (left) and proteomics (right). Each point represents one gene; genes passing the FDR threshold (q < 0.10, Benjamini-Hochberg) are labelled. Differential expression was computed using limma (robust lmFit and eBayes). x-axis: log_2_ fold change between subgroup and remaining cell lines; y-axis: −log_10_(FDR-adjusted p-value). Abbreviations: RMSE, root mean square error.

To assess failure modes in the generalisation of DMMs, we performed leave-one-out cross-validation (LOOCV) across all cell lines in the training set and quantified the generalisation gap (ΔRMSE; Fig. 4B). ΔRMSE was defined as the difference between the prediction RMSE when a given cell line was held out for validation and its average RMSE when included in the training set. This analysis revealed two groups of cell lines with distinct patterns of elevated generalisation gap: *gap group G1* (orange; HCC2218) with high p-ERBB2 gap under +MEKi conditions and *gap group G2* (green; CAL51, MDAMB157, EVSAT, UACC893, MACLS2, and BT474) with high p-p90^rsk^ gap across conditions. Notably, cell lines with high error under +MEKi conditions for p-ERK and p-p90^rsk^ (purple, *E3*) also often had high gap for the same marker/condition combinations.

Poor model predictions for HCC2218, the only member of *G1* (orange), were previously reported for regression models in the context of the DREAM challenge that introduced the dataset (Gabor et al., 2021), pointing to a lack of predictive features rather than missing mechanistic knowledge as the underlying cause. We therefor turned to mutational information in the Cell Line Encyclopedia (CCLE) (Barretina et al., 2012), the Catalog Of Somatic Mutations In Cancer (COSMIC) (Forbes et al., 2015) and whole exome sequencing (WES) for the employed cell line panel (Marcotte et al., 2016). A MET exon-14 deletion event, which impairs receptor degradation (Kong-Beltran et al., 2006), is annotated for this cell line alone within the panel (Fig. 4C) in CCLE and COSMIC. Moreover WES identified a MET^N375S^ mutation for HCC2218 and UACC893, which was likely filtered out in COSMIC and CCLE as MET^N375S^ is a germline mutation (Tengs et al., 2006) with a population frequency of about 10%. However, MET^N375S^ has also been reported to induce heterodimerisation of ERBB2 with MET and ligand-independent activation (Kong et al., 2020). We hypothesise that these MET alterations drive the observed p-ERBB2 gap pattern, though missing p-ERBB2 data for UACC893 precluded a direct test.

Similarly, DU4475, the only member of *E2* (green), exclusively harbours a BRAF^V600E^ mutation. We note that a BRAF^V600E^ mutation has also been reported for MDAMB361 in the CCLE (Barretina et al., 2012), but not in COSMIC (Forbes et al., 2015) or in WES (Marcotte et al., 2016), possibly reflecting differences in variant calling pipelines (Hudson et al., 2014) or cell line divergence. However, as the data-generating study (Tognetti et al., 2021) employed the same cell line branches as in the WES data, and BRAF^V600E^ can affect MEKi potency (Fröhlich et al., 2023; Gerosa et al., 2020), we found it most likely that BRAF^V600E^ was only present in DU4475, explaining the high p-ERK MEKi gap in this cell line, which, though not directly validated here, is the most parsimonious explanation.

Surprisingly, this analysis did not surface MDAMB231, which carries activating KRAS and BRAF mutations (KRAS^G13D^ and BRAF^G464V^). This cell line was unobtrusive in both error and gap, likely because these mutations raise baseline pathway activity without altering the MEKi response, a constitutive shift that can be captured by the CyTOF-based features. For the remaining error groups *E1* and *E4*, we identified no candidate signalling-altering variant in CCLE, COSMIC, or WES. These discrepancies may instead reflect data-quality limitations, undiscovered functional variants or mechanisms outside the modelled pathway, which we did not pursue further.

Because variant calling in cell lines is itself error-prone and the functional consequence of a called variant is often uncertain, this points to a complementary role for DMMs: model failure provides a functional readout that flags variants reshaping signalling responses, and, when cross-referenced with prior literature, builds confidence in both their presence and their functional relevance.

### Model failure identifies candidate rare signalling programmes driving MEK inhibitor resistance and ERK to p90^rsk^ gain

Encouraged that model failures pinpointed interpretable, mutation-driven biology in individual cell lines, we next analysed the larger error and gap subgroups E3 and G2, where recurrence across cell lines allows associations to be supported statistically.

For high error group *E3* (purple), we observed elevated p-ERK under +MEKi treatment, indicative of on-target MEK inhibitor resistance (Fig. 4D, top) and consistent with previous reports (Aksamitiene et al., 2010; Tognetti et al., 2021). The mechanistic model trained on individual cell lines (MM, dotted line), which supports MEK-independent ERK activation proposed in these reports, captured elevated p-ERK in this group, whereas the DMM (solid line) showed only modest increases, pointing to a lack of informative features. To identify more informative features, we performed differential expression analysis (Fig. 4E, top panels), excluding CAL148 which harbours a MAP2K1^E203K^ mutation conferring MEKi resistance (Mizuno et al., 2023). This yielded four statistically significant associations with transcript levels (FDR<10%, Benjamini–Hochberg correction): CCDC30, ZNF175, SLC6A1 and PRKAA2. Of these, PRKAA2 (log_2_FC=-2.18, q=0.06) is the only gene with known direct interactions with ERK signalling. PRKAA2 encodes the α2 catalytic subunit of AMPK; AMPK phosphorylates BRAF at Ser729, its C-terminal 14-3-3 binding site, promoting 14-3-3 recruitment and disrupting the BRAF–KSR1 association, thereby dampening MEK phosphorylation (Shen et al., 2013). Both 14-3-3 and KSR1 have been recognised as important determinants of MEK-inhibitor engagement and action (Khan et al., 2020; Marsiglia et al., 2024). Notably, AMPKα1 activation by phosphorylation at Thr172 is inhibited by Akt mediated phosphorylation at Ser485, while the equivalent Ser491 site in AMPKα2 is not phosphorylated by AKT, which may help explain why MEKi resistance in these cell lines was observed to be AKT-dependent (Aksamitiene et al., 2010).

For the high gap group *G2* (green, bottom panels, Fig. 4D–E), DMMs captured lower p-p90^rsk^ levels more accurately when cell lines were included in the training set (solid light green line, Fig. 4D bottom) than when held out for validation (dashed light green line), again suggesting a lack of predictive features rather than a shortcoming of the mechanistic description. Differential gene expression analysis (Fig. 4E, bottom panels) identified 6 transcripts and 22 proteins with statistically significant associations, among which 11 (see Table S5) are involved in cytoskeletal dynamics (GSN, STXBP2, TAGLN2, KIFAP3, REEP3, TTC21A), endo/exocytosis (DOC2A, HTATIP2, PRAF2) or calcium sensing/dynamics (GSN, DOC2A, STXBP2, SUMO1, S100A11). Notably, both transgelin 2 (TAGLN2) and gelsolin (GSN) couple ERK signalling to the actin cytoskeleton (Morley et al., 2007; Sun et al., 2018). The similarity in the affected model readouts and in the gene programmes enriched among the associated proteomic and transcriptomic features suggested that the high-gap group reflects a failure of CyTOF-based DMMs to predict the *P1* heterogeneity axis, which we confirmed by finding statistically significant differences in estimated parameter values between the training and test sets (Fig. S8).

To validate the identified gene programmes, we scanned the high-gap group for mutations that were classified as oncogenic in OncoKB (Chakravarty et al., 2017) or pathogenic in ClinVar (Landrum et al., 2016) and identified a cluster of alterations in Rho GTPases exclusively in three of the six high-gap group cell lines (Fig. 4C): RAC1^N39S^ in CAL51, RAC1^R9S^ in MDAMB157, and RHOA^G17E^ in BT474. As Rho GTPases are key regulators of the cytoskeleton (Etienne-Manneville and Hall, 2002; Hall, 1998) this, together with the reported role of p90^rsk^ in cell motility and cytoskeletal remodelling (Doehn et al., 2009; Sulzmaier and Ramos, 2013) suggest that the heterogeneity axis *P1* likely does not just reflect alterations in Ca^2+^/p38 signalling but also cytoskeletal remodelling. However, we observed no correlation with cell invasion assays (Neve et al., 2006) or association with morphology classes (Kenny et al., 2007) (Fig. S9) arguing against a canonical pro-migratory phenotype.

Together, these analyses establish that the failure modes of DMMs are a means of discovering structure in the data: recurrent error and gap signatures, and the molecular features associated with them, discover patterns that would otherwise go unnoticed. These patterns can corroborate known associations and point to novel candidate biology, here a putative AMPK–BRAF resistance axis and a cytoskeletal/secretory *P1* axis, though the underlying mechanisms remain to be experimentally validated.

### In silico integration triages mechanistic hypotheses and points to architecture as the limiting factor

Throughout our model analyses we identified several mechanistic hypotheses, such as ligand-driven baseline ERBB2 phosphorylation driving the *P2* heterogeneity axis, BRAF^V600E^ mutations altering MEKi response, loss of PRKAA2 expression driving MEKi resistance, or involvement of cytoskeletal remodelling in the *P1* heterogeneity axis. Although most of these hypotheses could be tested experimentally, reproducing the exact setup and validating predictions across a large cell-line panel would require substantial effort. Instead, to triage these mechanisms, we filtered the hypotheses through model predictions following mechanistic or feature-set integration.

For the hypothesis of ligand-driven ERBB2 basal phosphorylation, we explored mechanistic integration by extending the mechanistic model to include activation of EGFR by BTC, EREG and TGFA, and of ERBB3 by NRG1 and NRG2 (see Methods Section 2.4). None of these integrations improved model predictions (Fig. 5A), suggesting either that paracrine signalling is not a major contributor to signalling heterogeneity, or that additional regulatory mechanisms, such as shedding by metalloproteases or receptor dimerisation, are needed to reproduce the effect on basal phosphorylation.

**Figure 5.**
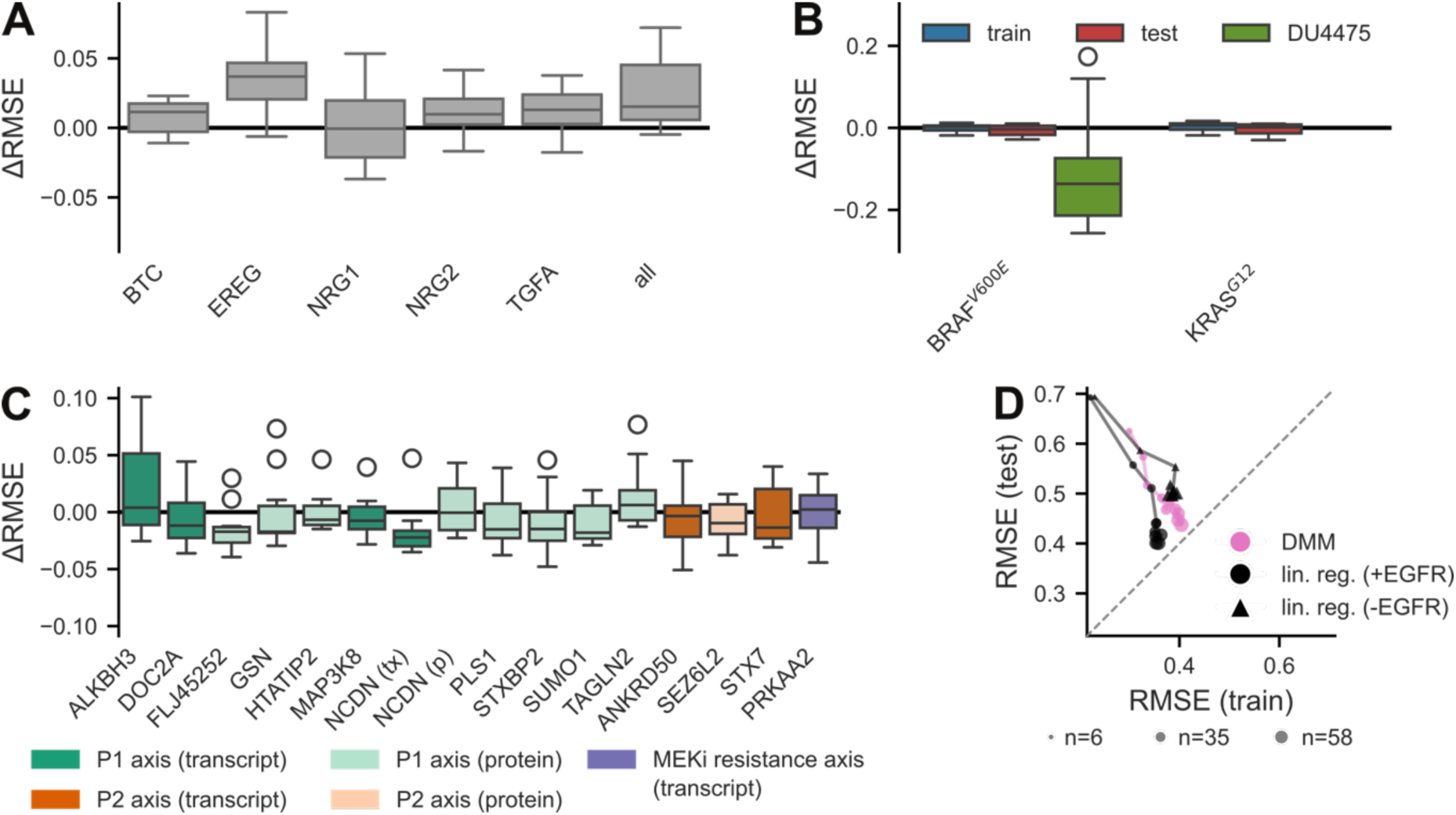
Mechanistic integration, feature augmentation, and training data scaling in deep mechanistic models. (**A**) ΔRMSE (RMSE after additional mechanistic integration of growth factor ligands (BTC, EREG, NRG1, NRG2, or TGFA—or all five simultaneously (all)) minus RMSE after mechanistic integration with EGFR transcript levels) evaluated on the held-out test set. **(B)** ΔRMSE upon mechanistic integration of oncogenic mutations BRAF^V600E^ or KRAS^G12^ evaluated on the training set (train), held-out test set (test), and the DU4475 cell line from the training data. **(C)** ΔRMSE upon single-marker feature augmentation with individual protein (p, light color) or gene-expression (tx, dark colour) measurements. Markers are coloured by heterogeneity or gap axis from Fig. 3,4: *P1* axis (green shades), *P2* axis (orange shades) and MEKi gap (purple shade). **(D)** Mean training RMSE versus mean test RMSE across seeds for DMM and linear regression (lin. reg.) with elastic net regularisation models trained on subsets of the available cell lines. Dot size reflects the number of cell lines used for training. Marker shape reflects employed model and feature set. For (A–C), boxes show the interquartile range (IQR); whiskers extend to 1.5 × IQR.

To capture the effect of the oncogenic driver mutations BRAF^V600E^ and KRAS^G13D/G12^, we performed mechanistic integration by adding a constitutively active MEK term, gated by the presence of either mutation (see Methods Section 2.4). For KRAS^G13D^ (Fig. 5B, right) we observed negligible change in RMSE (ΔRMSE=RMSE_integrated_-RMSE_baseline,_ negative values indicate reduced error) on the training (blue, ΔRMSE=0.002) or test (red, ΔRMSE=-0.003) set, consistent with our earlier observation that this mutation was not associated with high error or a large generalisation gap. Surprisingly, mechanistic integration of BRAF^V600E^ (Fig. 5B, left) likewise had negligible impact on the training (blue, ΔRMSE=-0.0005) or test (red, ΔRMSE=-0.005) set. At the level of individual cell lines, however, it significantly reduced the p-ERK +MEKi error for DU4475 (ΔRMSE=-0.1), suggesting that, given the rarity of the mutation, its net effect across the cell-line panel is too diluted to be detectable. Moreover, the data-driven component cannot account for the BRAF^V600E^-driven contribution to p-ERK in the input features, which may bias the learnt latent representation.

Unlike single-cell-line effects, which are diluted across the panel and best assessed per line, heterogeneity programmes recurring across many cell lines should improve panel-level predictions if genuine. However, many of the respectively identified genes are not part of the EGFR/MAPK signalling pathway, making them incompatible with mechanistic integration. Instead, we performed feature augmentation for transcripts (orange) and proteins (teal) with established links to EGFR/MAPK signalling, the Ca^2+^/p38 axis, the cytoskeleton or endo-/exocytosis. Candidates were required to be the strongest positive and negative correlates for with *P1* (ERK to p90^rsk^ signal gain; PLS1, NCDN, MAP3K8) or *P2* (baseline ERBB2 activation; ANKRD50, SEZ6L2, STX7) heterogeneity axes for any modality, or among top-5 differentially expressed genes for high error group *E3* (MEKi resistance; PRKAA2) or high gap group *G2* (ERK to p90^rsk^ signal gain; HTATIP2, TAGLN2, GSN, SUMO1, FLJ45252, STXBP2, ALKBH3, DOC2A) in any modality, ranked by multiple-testing corrected p-values (Fig. 5C). Collectively, feature augmentations improved performance on the test-set in terms of median ΔRMSE (13/16, p=0.02 sign test), but none of the improvements were individually statistically significant after multiple testing correction. PRKAA2 was one of the three augmentations that did not reduce test error (median ΔRMSE = 0.002), providing no independent support and tempering the hypothesis advanced above. Individually, FLJ45252 yielded the largest reduction in error (median ΔRMSE = –0.017). Notably, FLJ45252 protein abundance was the only feature that we correlated with the *P1* axis and at the same time was differentially expressed in the associated *G2* group. Even though UniProt (The UniProt Consortium, 2015) lists FLJ45252 as uncharacterised protein, it has been proposed to be a putative isoform of AP2-associated protein kinase 1 (AAK1) (Huang et al., 2018), a clathrin-stimulated kinase that phosphorylates the AP-2 μ2 subunit (AP2M1) to drive clathrin-mediated endocytosis. Notably, AAK1 has also been linked to actin cytoskeletal remodelling (Chen et al., 2022), bolstering the association of the *P1* heterogeneity axis with endocytic and cytoskeletal remodelling.

Encouraged by the observation that mechanistic integration and feature augmentation can improve model performance on both training and test sets, we sought to identify the most promising avenues for further improvement. To this end, we ablated training-set size and evaluated training and test RMSE (Fig. 5D), comparing DMMs (pink) and linear regression with elastic net regularisation (black) trained on the same CyTOF features, with augmentation by EGFR transcript levels (circles) or without (triangles). For both models, test RMSE showed diminishing returns beyond 30 training cell lines, alongside a low generalisation gap. Linear regression reached a lower test RMSE than the DMM, but only when EGFR transcripts were included as features, and chiefly by attaining a lower training RMSE that carried through to the test set without a widening gap. This indicates that the predictive signal is learnable from the available data, and that the DMM’s higher error reflects the limited capacity imposed by its mechanistic structure rather than a shortage of data. The comparison is nonetheless asymmetric: the DMM’s parameters map directly onto pathway processes, whereas the linear model distributes each feature’s contribution across time points, observables, and perturbation conditions, so its coefficients do not resolve into an interpretable, mechanism-level account. We thus view this predictive gap as a price paid for a representation that supports the mechanistic and failure-mode analyses above. Moreover, it suggests that the most promising route to further gains lies in mechanistic model architecture, driven by a better understanding of the underlying biology rather than in more data.

Together, these findings demonstrate that both mechanistic integration and model-guided feature augmentation can improve model performance. Moreover, where either approach fails to yield such improvements, it remains informative, acting as a filter that flags hypotheses not worth costly experimental follow-up, as with PRKAA2, or as a means of refining a mechanistic hypothesis in advance, as with baseline ERBB2 phosphorylation. At the same time, the diminishing returns observed with increasing training-set size suggest that the largest remaining gains lie in model architecture, marking this as the priority for future development.

## Discussion

In this manuscript, we introduce Deep Mechanistic Models (DMMs), a novel framework that blends machine learning with mechanistic mathematical modelling. We show that, in contrast to purely statistical models, the mechanistic component provides a rich scaffold for downstream interpretation: it allowed us to identify three axes of heterogeneity: the two major axes recovered from sensitivity analysis of the inferred parameter deviations, baseline ERBB2 phosphorylation and signal transduction from p-ERK to p-p90RSK (Fig. 3), together with EGF sensitivity, which we uncovered via inspection of the learnt latent representation (Fig. 2B) and mechanistic integration EGFR levels (Fig. 2E). All three indicate that heterogeneity manifests primarily at the level of pathway inputs and outputs. In contrast to purely mechanistic models, DMMs can also integrate variation originating outside the pathway under investigation; this suggested that, although heterogeneity manifests at the pathway’s inputs and outputs, much of this heterogeneity is associated with pathway-extrinsic factors such as Ca^2+^/p38 signalling or cytoskeletal remodelling rather than core MAPK pathway members, which appear to have little predictive value, confirming previous reports (Shi et al., 2016). Finally, by mechanistically integrating EGFR transcript levels, we show that the mechanistic component shapes the latent representation learnt by the machine learning component. We exploit this feature of the model architecture to show that a molecular lens—implemented as a CyTOF-based DMM with mechanistic integration—and a systems lens— implemented as a multimodal DMM without mechanistic integration—are equivalent in predictive power.

DMMs critically rely on the mechanistic model. We deliberately employed a simple, coarse-grained description of EGFR and MAPK signalling to keep computational runtime manageable, reduce potential bias from model misspecification and enable the machine learning component to learn context-specific nuances. Overall, we find that this model was sufficiently expressive to describe most of the experimentally observed dynamics, except for transient pathway activation in the context of +RTKi conditions. Still, although the mechanistic model affords greater interpretative depth, we must establish that model-suggested patterns are genuine rather than modelling artefacts: confirming the identified heterogeneity axes and the model’s failure modes in experimental data renders our findings and their interpretation independent of the model employed.

CyTOF-based features generally yield the best predictive performance, and we find surprisingly little statistical signal when associating heterogeneity axes or model failure modes with transcriptomic or proteomic data. It would be easy to dismiss this as a data artefact: Because the prediction target is itself a CyTOF measurement, CyTOF-based features enjoy a natural advantage; the transcriptomic and proteomic data, by contrast, were not collected under the same conditions, or even by the same laboratories, and may simply be too noisy to be informative. However, the mechanistic integration of EGFR transcript values, collected five years earlier in a different laboratory, yielded the single largest improvement in model performance, confirming that these datasets do carry meaningful signal. The observation of individual mutations, such as MAP2K1^E203K^, that are consistent with the identified signalling phenotypes—here, MEKi resistance—suggest the possibility that each cell line has its own idiosyncratic molecular implementation of a given signalling phenotype, casting doubt on whether a purely statistical approach can extract transferable structure. Here, DMMs offer a powerful means of accounting for such rare alterations through mechanistic integration, as we demonstrate for the BRAF^V600E^ mutation, although this requires a reliable mechanistic understanding of the alteration’s mode of action. The strong performance of CyTOF-based DMMs, however, suggests that these idiosyncratic alterations can also converge on emergent cellular programmes, some of which are reflected in phosphorylation levels. The machine-learning component should in principle be able to learn these programmes from transcriptomic and proteomic data, but limited data availability probably constrained our ability to learn them reliably, forcing us to resort to aggressive feature selection prior to DMM training. The alternative to more samples of a specific data type could be additional data modalities: our difficulty in reliably predicting the *P1* heterogeneity axis, which is putatively governed by cytoskeletal remodelling, would most plausibly be addressed by imaging that captures morphology, which DMMs could accommodate naturally, since the machine-learning component could easily include an additional convolutional encoder head.

Agnostic to both input modality and mechanistic model, DMMs offer a general way to reconcile the molecular and systems-level descriptions of signalling that have largely developed apart, making the two mutually legible on the same data rather than forcing a choice between them. To our knowledge this is the first framework to couple representation learning to a mechanistic signalling model such that either view can inform the other. Extending it across further modalities and pathways would let these two views be held together throughout the signalling networks that shape cell fate.

## Acknowledgements

We thank the whole Dynamics of Living Systems lab and Colin Ratcliffe for helpful discussions and feedback on the manuscript.

## Material and Methods

### 1. Software and computational modules

The framework is implemented in Python and builds on the following open-source packages. The three core modules are **AMICI** (Fröhlich et al., 2021) (model import, simulation and sensitivity analysis), **JAX** (Bradbury et al., 2018) (automatic differentiation and just-in-time compilation of the full hybrid objective), and **pyPESTO** (Schälte et al., 2023) (parameter-estimation problem assembly and the bridge between AMICI and JAX).

**Supplementary Table S1.** Software modules used in the deep mechanistic modelling pipeline.

| Module | Role in the pipeline | Version |
| --- | --- | --- |
| PySB (Lopez et al., 2013) | Programmatic, rule-based construction of the mechanistic model; export to SBML / flat-PySB. | 1.17.0 |
| BioNetGen (Blinov et al., 2004) | SBML representation and reaction-network generation from PySB rules. | 2.8.5 |
| AMICI (Fröhlich et al., 2021) | Code generation (C++) for the ODE right-hand side and sensitivities; model compilation and simulation. | 0.33.0 |
| PEtab (Schmiester et al., 2021) | Standardised specification of conditions, measurements, observables and parameters. | 0.5.0 |
| pyPESTO (Schälte et al., 2023) | Objective construction (AmiciObjective) and JAX bridge (JaxObjective). | 0.5.6 |
| JAX (Bradbury et al., 2018) | Reverse-mode autodiff, vmap over cell lines, and JIT compilation of the training loss. | 0.6.2 |
| Equinox (Kidger and Garcia, 2021) | Neural-network module definitions (encoder / inflater / decoder). | 0.13.0 |
| optax | Adam optimiser and learning-rate schedule. | 0.2.5 |
| scikit-learn | Imputation, random-forest RFE, baseline regressors, PCA, scaling, Gaussian mixture model. | 1.8.0 |
| SciPy | Correlation analysis and multiple-testing correction. | 1.17.1 |

### 2. Mechanistic model formulation

#### 2.1 Programmatic model construction

The mechanistic model is assembled programmatically with PySB (dmm/mechanistic_model.py). Each signalling protein is represented by a PySB *Monomer* carrying phosphosite and compartment states. Protein synthesis and degradation can be added as mass-action rules (`add_monomer_synth_deg`), and every regulated (de)phosphorylation, endocytosis or degradation event is added as an activation/deactivation rule (`add_activation`). The biological wiring, which species activate or inhibit each site, is declared in a single pathway specification (`cytof/pathways.py`) as a tuple of (activators, inhibitors) for each target site. The completed PySB model is exported both to SBML and to a flat-PySB Python file via `pysb.export.export(model, “pysb_flat”)`; SBML is the input to AMICI.

The model is modular: EGFR and core MAPK reactions form the base model (EGFR_MAPK), with `logobs` suffix denoting log-transformed observables to account for asinh transformation. Modifier flags toggle additional inputs such as activating mutations (`m_KRAS`, `m_BRAF`) and additional growth-factor ligands.

#### 2.2 Functional form of the mechanistic integration

The central modelling choice is how multiple activators and inhibitors of a single site are integrated into one reaction rate. For each regulated site, modulator observables enter through log-scale weights *w* (parameter class kw) and a baseline term *b_act_* (bact). The default form is a rational (saturating in the inhibitors) law:

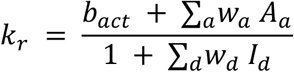

where *A_a_* and *I_d_* are activator and inhibitor (deactivator) observables (see Table S3). The realised reaction rate scales this dimensionless factor by the site’s reference catalytic/degradation constant:

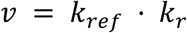

Where *k_ref_* is the reference *k_cat_* (catalytic: phosphorylation/endocytosis) or *k_deg_* (degradation). Endocytosis is modelled with a three-step delay chain and requires prior phosphorylation of the receptor; degradation and membrane reactions are gated on compartment state.

#### 2.3 Parameters, scales and observables

All estimated parameters are bounded and sampled on a log₁₀ scale. Observables are modelled as log-transformed, affinely scaled readouts, *y* = log(*s* · *x_obs_* + *o*), with observable-specific scale *s* and offset *o*.

**Supplementary Table S2.** Parameter classes, optimisation scales and bounds (cytof/problem.py).

| Parameter class | Scale | Lower | Upper | Description |
| --- | --- | --- | --- | --- |
| $k_{deg}$ | $\log_{10}$ | -4 | 0 | Degradation rate constants |
| eq | $\log_{10}$ | -4 | 4 | Total protein levels / equilibrium concentrations |
| $k_{cat}$ | $\log_{10}$ | -3 | 3 | Catalytic (phosphorylation / transport) constants |
| scale | $\log_{10}$ | -2 | 3 | Observable scaling factor $s$ |
| offset | $\log_{10}$ | -4 | 3 | Observable offset $o$ |
| kw | $\log_{10}$ | -3 | 3 | Modulator (activator / inhibitor) weights $w$ |
| bact | $\log_{10}$ | -5 | 3 | Baseline activation |

#### 2.4 Reaction network

Supplementary Table S3. enumerates the regulated sites of the mechanistic model with their activators and inhibitors. Throughout, the active receptor tyrosine kinases are `active_rtks` = (`EGFR Y1173_p`, `ERBB2 Y1248_p`, `ERBB3 Y1289_p`). Pharmacological inputs are the receptor and MEK inhibitors (`iEGFR_0`, `iMEK_0`), and stimuli are EGF and serum (`EGF_0`, `serum_0`).

**Supplementary Table S3.**
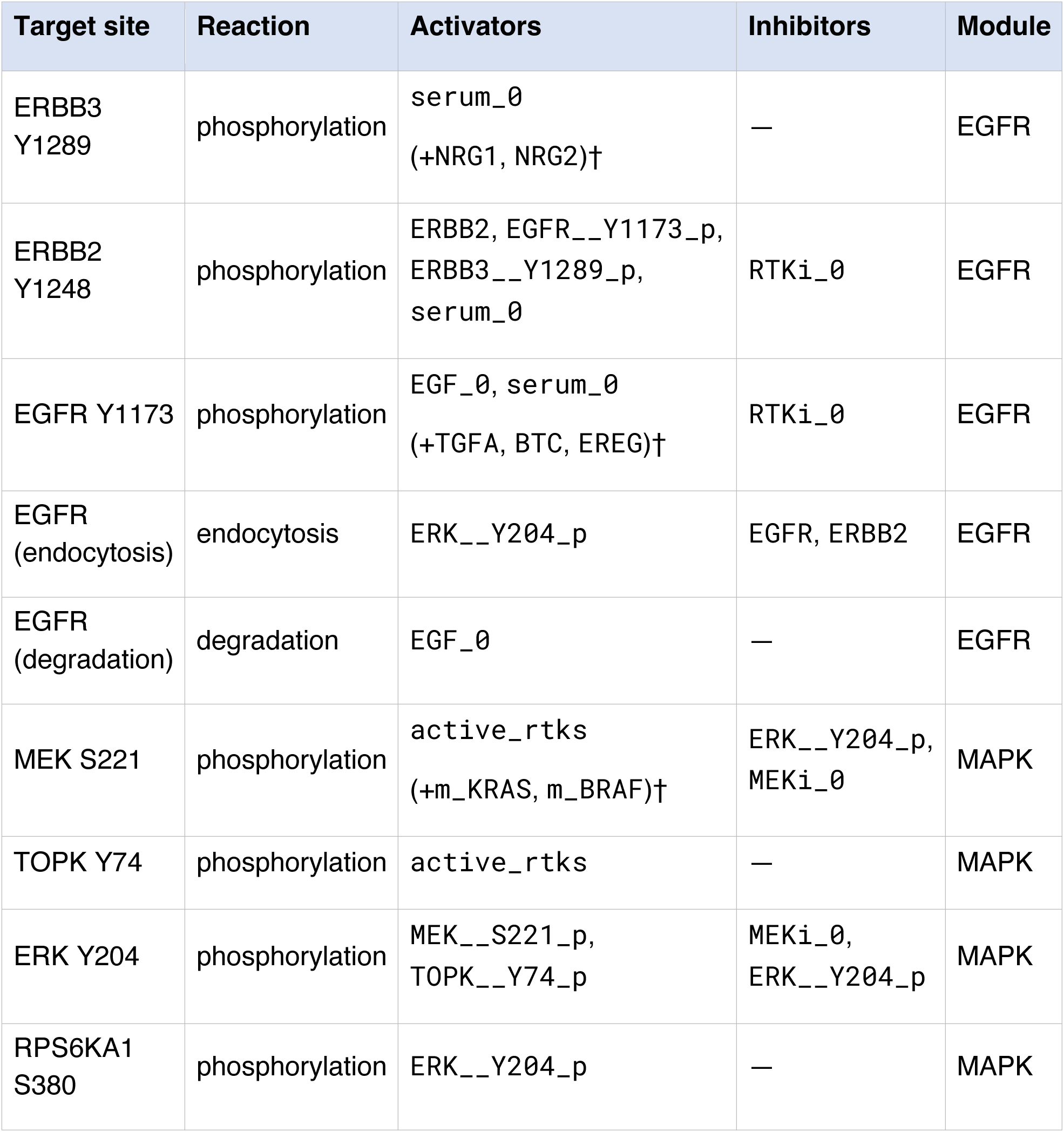
Regulated sites, reaction types, activators and inhibitors (cytof/pathways.py). † Inputs in parentheses are included only when the corresponding mutation/ligand modifier flag is set. ‡ Endocytosis uses a three-step delay chain and requires Y1173 phosphorylation. Basal synthesis and degradation reactions are only added for EGFR.

### 3. Mechanistic model simulation

The PySB/SBML model is imported into a PEtab problem (dmm/petab_subproblem.py) consisting of condition, measurement, observable and parameter tables. A pyPESTO PetabImporter then generates and compiles the AMICI model, which produces optimised C++ code for the ODE right-hand side and its sensitivities and links against the SUNDIALS CVODES (Serban and Hindmarsh, 2005) integrator (compile_model.py). For training, the resulting `AmiciObjective` is wrapped in a custom χ² objective and exposed to JAX through pyPESTO’s `JaxObjective`, providing both the objective value (sensitivity order 0) and its gradient (sensitivity order 1) to the JAX training loop.

#### 3.1 Steady states and pre-simulation

Each condition is first integrated to steady state (pre-equilibration) with all perturbations (`EGF_0`, `serum_0`, `iMEK_0`, `iEGFR_0`) set to zero. The steady state is found by integration, and steady-state sensitivities are computed with Newton’s method. Pre-simulation implements the experimental design: for perturbation conditions, the stimulus parameters `EGF_0` and `serum_0` are kept to zero while corresponding drug-parameters (`iMEK_0` or `iEGFR_0`) are set to 1 (arbitrary units) during a pre-simulation phase of duration *t*_presim_ = 15 minutes. For simulation itself, EGF_0 and serum_0 are also set to 1 (arbitrary units). State variables are not reinitialised between pre-equilibration and pre-simulation, or between pre-simulation and simulation, so protein levels (e.g. EGFR) carry over. For “full” conditions, only serum_0 is set to 1 during pre-equilibration with no pre-simulation or simulation phases.

#### 3.2 Integration and sensitivity settings

First-order sensitivities are used for gradient-based training. No explicit sensitivity method is set, so AMICI’s default forward sensitivity analysis is applied during dynamic phases, with Newton-based steady-state sensitivities as above. Solver tolerances are:

- absolute / relative tolerance = 10⁻¹⁰
- steady-state absolute / relative tolerance = 10⁻⁶
- maximum integration steps = 2×10⁴, maximum Newton steps = 100
- Newton-step steady-state convergence check enabled.

### 4. Neural network architecture

The neural network is implemented in Equinox and comprises three sub-modules (dmm/dmm_autoencoder_eqx.py, dmm/deepcomponent_eqx.py): an encoder mapping omics features to a latent embedding of dimension *n*_hidden_; an inflater mapping the latent embedding to the cell-line-specific kinetic-parameter deviations *Δ*; and an optional decoder reconstructing the input features (instantiated only when the reconstruction-loss weight is non-zero). All cell lines are processed in parallel with `jax.vmap`.

#### 4.1 Parametrisable depth and width

The architecture is fully parametrised by its `depth` (number of hidden layers, default 0) and latent dimension (n_hidden_, default 4). Hidden-layer widths follow a geometric schedule (dmm/model_utils.py) with respect to n_hidden_ (default: 4). Inflater and decoder grow from n_hidden_, with successive layers being multiplied by the structure factor m (`nn_structure_multipler`=2) up to their output dimension. The encoder, instead, reverses this schedule: it can first expand to a maximum width equal to 3 x (number of input features), and then contracts to n_hidden_. The maximum widths of encoder/decoder are coupled, while that of the inflater is capped at the number of kinetic-parameter deviations. To bridge the final gap between the last geometric progression and the respective maximum width of each module, the structure factor is raised using a smoothing rule up to a maximum of 5. The inflater’s output layer carries no bias, preserving the scaling of the predicted deviations.

#### 4.2 Activations, regularising layers and multimodal inputs

- **Dropout:** applied to the encoder only, rate 0.1 by default; no batch or layer normalisation; layer biases off by default.
- **Multimodal (“multi-headed”) setup:** when enabled, a separate encoder (and decoder) is instantiated per input modality (e.g. proteomics, transcriptomics, CyTOF-derived features); the per-modality latent embeddings are mean-pooled into a single shared latent embedding that a single inflater consumes, and per-modality decoder outputs are concatenated (dmm/two_headed_deep_autoencoder_eqx.py).

#### 4.3 Coupling to the mechanistic model

The inflater output, i.e. the cell-line specific parameter deviations *Δ*(*Φ*) (with network parameters *Φ*) is added to the global kinetic parameters *Θ* so that cell line *c* is simulated with parameters *Θ + Δ*_c_. A binary sparsity mask multiplies *Δ* so that only parameters identified as genuinely cell-line-variable are allowed to deviate (Section 6).

### 5. Feature selection and preprocessing

Input omics features were filtered and reduced before training (select_features.py, dmm/feature_selection.py) in the following order:

1. **Missingness filter:** features missing in more than 30% of samples were removed
2. **Imputation:** remaining missing values were imputed with k-nearest-neighbours imputation (`KNNImputer`, `k` = 5).
3. **Low-expression filter:** the lowest 20% of features by mean expression were removed.
4. **High-variance selection:** the top *N* features by variance were retained (`get_hvg`, default `top_n` = 500; a fractional `top_n` argument instead keeps a top percentile).
5. **Recursive feature elimination (RFE):** starting from the high-variance set, a random-forest regressor (`RandomForestRegressor(random_state=42, max_features=0.80)`) was fit within a standardising pipeline; at each iteration the bottom ∼20% of features by importance were dropped (at least one feature per step) until further reduction by the 0.80 factor would fall below the target size, after which a final fit selected exactly the target number of features. Feature importances used permutation importance (`n_repeats`=10, `random_state`=42) when ≤ 500 features remained, and the faster impurity-based importances otherwise.

Selection was reproducible (fixed random_state = 42); output targets were mean-centred before RFE.

### 6. Model training

Training minimises a composite objective over the global kinetic parameters *Θ* and the network parameters *Φ* jointly, using the Adam optimiser (`optax.adam`) with a constant learning rate of 0.01. Optimisation runs for 500 epochs in full-batch mode (all cell lines vmapped together), with a fresh dropout key per epoch.

#### 6.1 Objective function

The training loss combines the mechanistic data-fit term with regularisers:

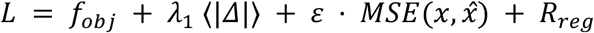

Here *f_obj_* is the AMICI/PEtab mechanistic data-fit term (weighted residual sum of squares / negative log-likelihood) evaluated at *Θ + Δ*; the second term is an L1 penalty on the magnitude of the predicted deviations (⟨·⟩ denotes the mean over deviations), which drives sparsity in the number of non-zero cell-line-specific parameters; the third is the optional feature-reconstruction mean-squared error; and *R_reg_* collects further optional penalties — L1 on encoder and inflater weights, an L2 penalty on the deviations, and orthogonality and encoder–decoder symmetry penalties. Default weights are *λ_1_* = 0.1 (L1 on deviations) and *ε* = 0.01 (reconstruction); all other regulariser weights default to 0, and the orthogonality penalty, when used, defaults to the L2 form.

#### 6.2 Sparsification: GMM selection and regularisation relief

Sparsity in the cell-line-specific parameters is enforced in two stages. During the first 200 epochs (`inflater_output_reg_epoch` = 200), the L1 penalty on the kinetic-parameter deviations, *Δ,* is active. At epoch 200, the per-parameter standard deviation of *Δ* across cell lines is computed, and a two-component Gaussian mixture model is fitted to their logarithms (`GaussianMixture(n_components=2, random_state=42`)). The threshold separating cell-line-variable from negligible parameters is taken as the geometric mean of the two component means:

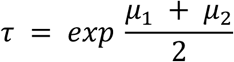

Parameters whose deviation standard deviation falls below *τ* are fixed at the global value by zeroing the corresponding entries of the binary sparsity mask. The L1 penalty on the inflater output is then disabled (“relieved”) for the remaining epochs, so that the surviving cell-line-specific parameters are estimated without shrinkage.

#### 6.3 Initialisation

- **Mechanistic parameters:** the global (median) kinetic parameters are initialised from a pre-trained “average” model fitted to the pooled data; one parameter set is sampled from this reference (dmm/initialisation.py).
- **Network parameters:** weights are initialised with variance scaling `(scale=0.1, mode=“fan_avg”, distribution=“uniform”)`; biases are zero and the inflater output layer is bias-free, so deviations start near zero.

### 7. Hyperparameter tuning

Hyperparameters were tuned by linear (one-dimensional) scans: each hyperparameter was varied along its own axis while the others were held at context-specific central values, and the central values were refined iteratively across rounds (generate_run_configs.py). Representative scan ranges were latent dimension n_hidden_ ∈ {1,…,8}, depth ∈ {0,…,4}, deviation L1 weight ∈ {0, 10⁻³, 10⁻², 10⁻¹, 1, 10}, and dropout ∈ {0, 0.05, 0.1, 0.2, 0.4}.

For each configuration, models were trained with 10 random seeds, and generalisation was assessed by leave-one-out cross-validation (LOOCV) over the five cell lines with the highest deviation from the mean (BT20, HCC1500, HCC2185, MCF7, UACC3199), with the remaining cell lines used for training. Performance was aggregated by averaging first over the random seeds and then over the held-out cell lines.

### 8. Baseline regression models

As baselines, the kinetic-parameter targets were predicted directly from omics features with linear models (regressor_training.py), each combined with the feature-selection schemes of Section 5:

- **Linear regression** (`LinearRegression`, multi-output).
- **Lasso** (MultiTaskLassoCV, cv=5, n_alphas=20).
- **Elastic net** (MultiTaskElasticNetCV, cv=5, n_alphas=20; the feature-selection variant uses l1_ratio ∈ {0.1, 0.5, 0.9}, n_alphas=25 with SelectFromModel).

All baselines used a common preprocessing pipeline: `StandardScaler` → `KNNImputer` → `regressor`. Missing values in the regression targets were KNN-imputed during training.

### 9. Statistical analysis

#### 9.1 Differential expression

Differential expression was performed with `limma` (Ritchie et al., 2015) called from Python via `rpy2` (figures_paper/embeddings.py). Features with more than 30% missingness were removed and the remainder KNN-imputed (`n_neighbors` = min(5, n − 1)); samples were split into two groups (median / positivity split of the variable of interest). A linear model was fitted (lm`Fit`, robust fitting when at least three samples per group), moderated with `eBayes(robust=TRUE)`, and ranked with `topTable` using Benjamini–Hochberg adjustment (`adjust.method=“BH”`).

#### 9.2 Correlation and multiple-testing correction

Pairwise associations (e.g. marker–parameter relationships) were quantified with Pearson and Spearman correlations (`scipy.stats.pearsonr / spearmanr`). Multiple-testing correction used the Benjamini–Hochberg false-discovery-rate procedure (`statsmodels.stats.multitest.multipletests(method=“fdr_bh”, alpha=0.05)`; `scipy.stats.false_discovery_control` was used as an equivalent alternative in places). Unless stated otherwise, significance was reported at an FDR of 0.05.

## Supplementary Information

**Supplementary Table S4.**
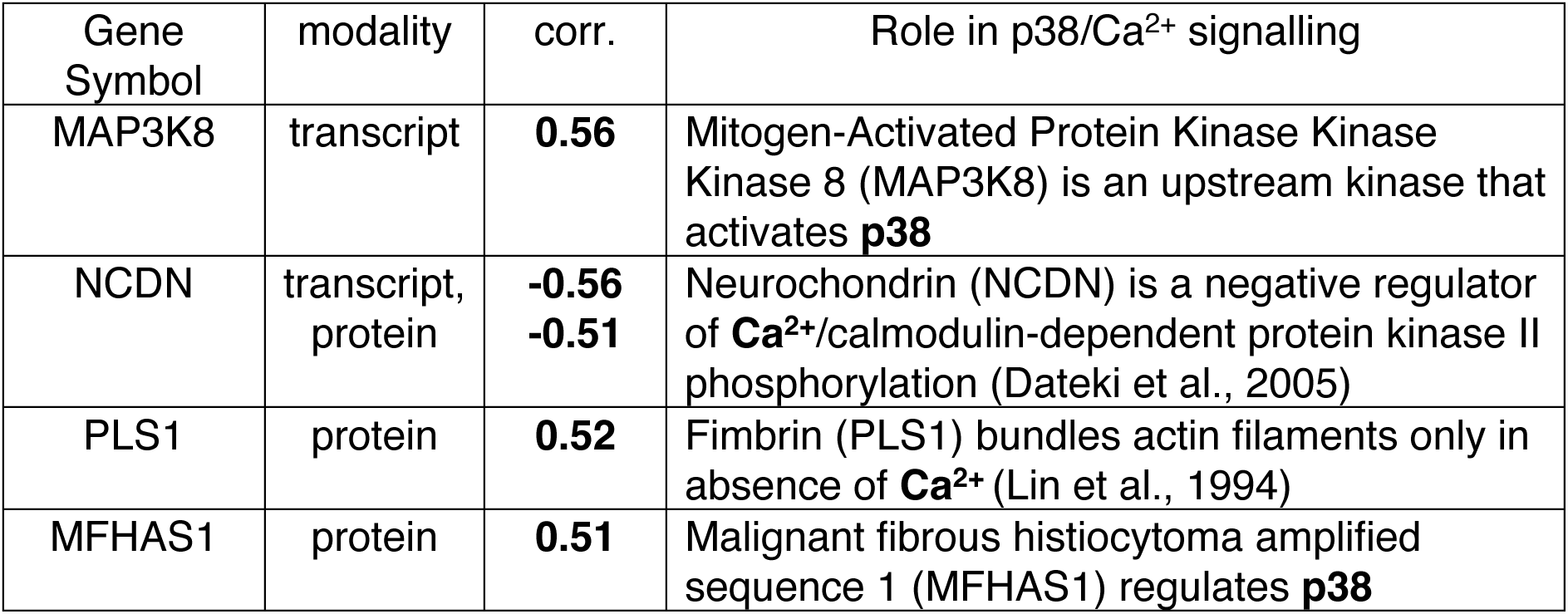
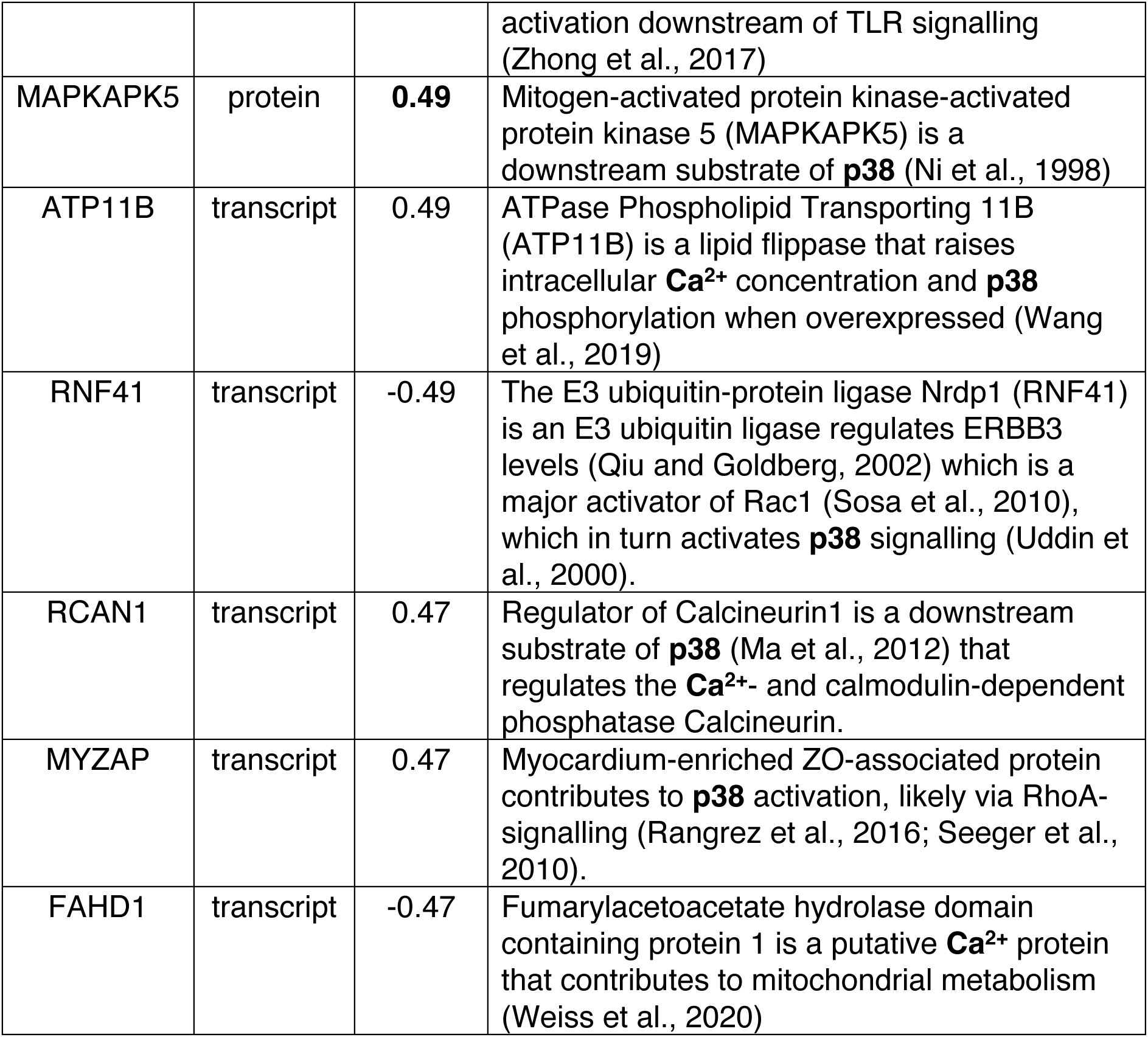
Gene mechanism analysis for *P1* heterogeneity axis.

**Supplementary Table S5.** Gene mechanism analysis for G2 gap group.

| Gene Symbol | modality | log<sub>2</sub>FC | Role in p38/Ca<sup>2+</sup> signalling, endo-/exocytosis or cytoskeleton |
| --- | --- | --- | --- |
| GSN | protein | **-2.32** | Ca<sup>2+</sup>-dependent actin filament severing and capping protein that drives dynamic reorganisation of the cortical actin **cytoskeleton**. (Serrander et al., 2000) |
| STXBP2 | protein | **-1.08** | Syntaxin-binding SM protein that is essential for SNARE-mediated regulated (including homotypic compound) **exocytosis** of secretory granules. (Gutierrez et al., 2018) |
| TAGLN2 | protein | **-1.63** | Actin-binding protein that stabilises actin filaments and thereby modulates **cytoskeletal** dynamics underlying migration and proliferation. (Dvorakova et al., 2014) |
| KIFAP3 | protein | **-0.70** | Non-motor accessory subunit of the heterotrimeric kinesin-2 (KIF3A/KIF3B/KAP3) complex that couples cargo to **microtubule**-based anterograde and ciliary transport. (Carpenter et al., 2015) |
| REEP3 | protein | **-0.97** | ER membrane-shaping/**microtubule**-linking protein that, redundantly with REEP4, clears ER from metaphase chromatin to maintain proper nuclear envelope architecture. (Schlaitz et al., 2013) |
| TTC21A | transcript | **-1.78** | IFT-A-associated TPR protein that mediates retrograde intraflagellar transport along the axonemal microtubule **cytoskeleton**. (Liu et al., 2019) |
| DOC2A | transcript | **-2.78** | Cytosolic tandem-C2-domain protein acting as a high-affinity **Ca<sup>2+</sup>** sensor that triggers spontaneous (Ca<sup>2+</sup>-dependent) vesicle **exocytosis** in competition with synaptotagmin-1 for SNARE binding. (Groffen et al., 2010) |
| HTATIP2 | protein | **-1.63** | Forms a complex with endophilin B1, ACSL4 and Rab5a that governs **endocytic** trafficking and lysosomal degradation of activated EGFR. (Zhang et al., 2011) |
| PRAF2 | protein | **+1.00** | PRAF2 is a four-TM, PRA1/Rab-acceptor-motif ER-membrane protein of the PRAF family (PRAF1/3) that regulates **endo-/exocytic** vesicle trafficking. (Borsics et al., 2010) |
| SUMO1 | protein | **+0.82** | Small ubiquitin-like modifier whose conjugation acts as a molecular switch promoting **Ca<sup>2+</sup>**-channel clustering and fast Ca<sup>2+</sup>-triggered vesicle **exocytosis**. (Girach et al., 2013) |
| S100A11 | protein | **-1.36** | **Ca<sup>2+</sup>**-binding S100 protein recruited with Annexin A2 upon injury-induced Ca<sup>2+</sup> influx to drive cortical F-**actin polymerisation** and plasma-membrane repair in motile cancer cells. (Jaiswal et al., 2014) |

**Figure S6.**
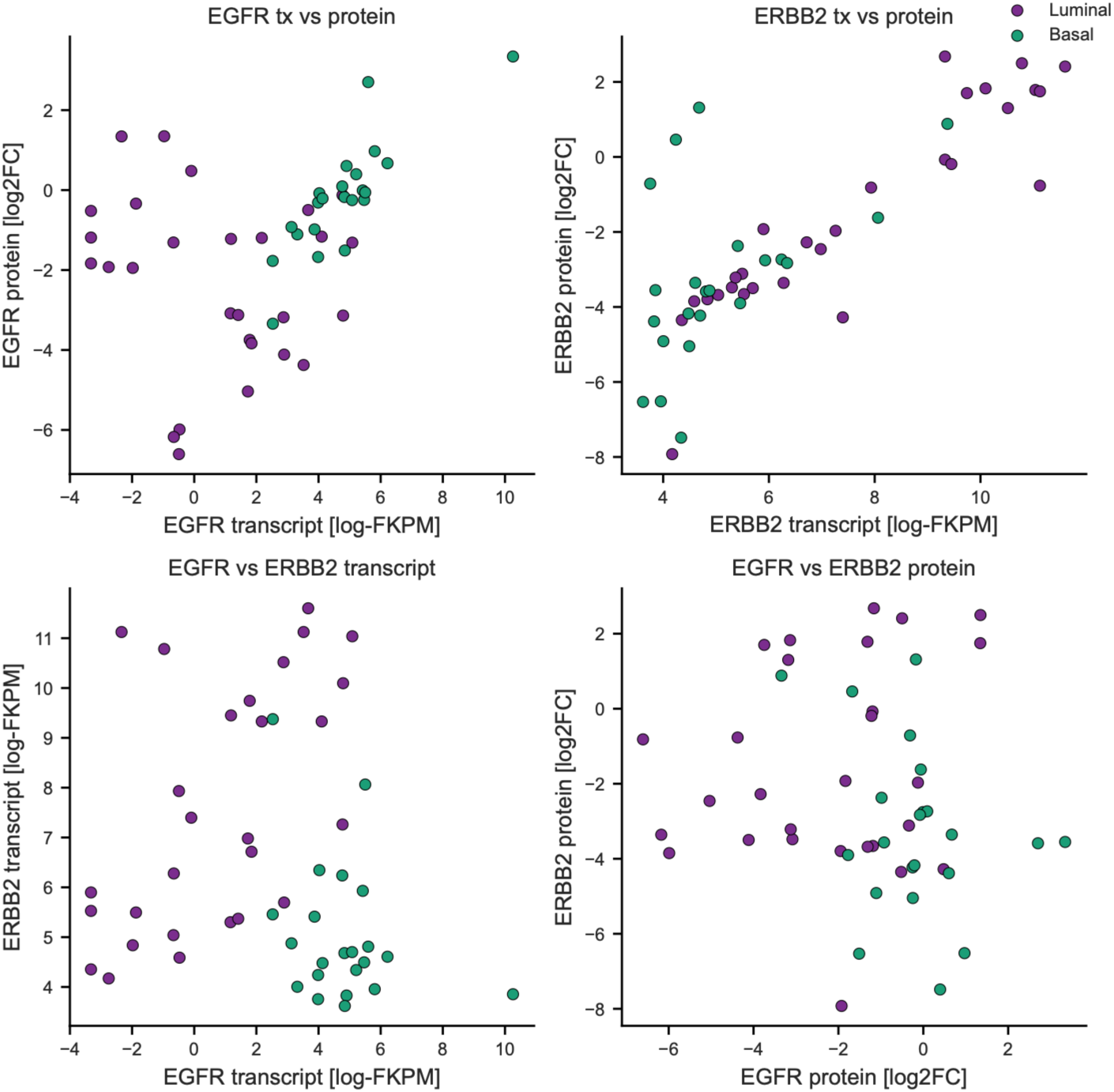
RTK transcript and protein expression across breast cancer cell lines. Pairwise scatter plots comparing EGFR and ERBB2 expression at the transcript (log-FKPM, RNA-seq) and protein (log₂FC relative to 5 normal cell lines, mass spectrometry) levels across 58 breast cancer cell lines (Luminal, n = 28; Basal, n = 22). Points are coloured by luminal (purple) and basal (teal) subtype.

**Figure S7.**
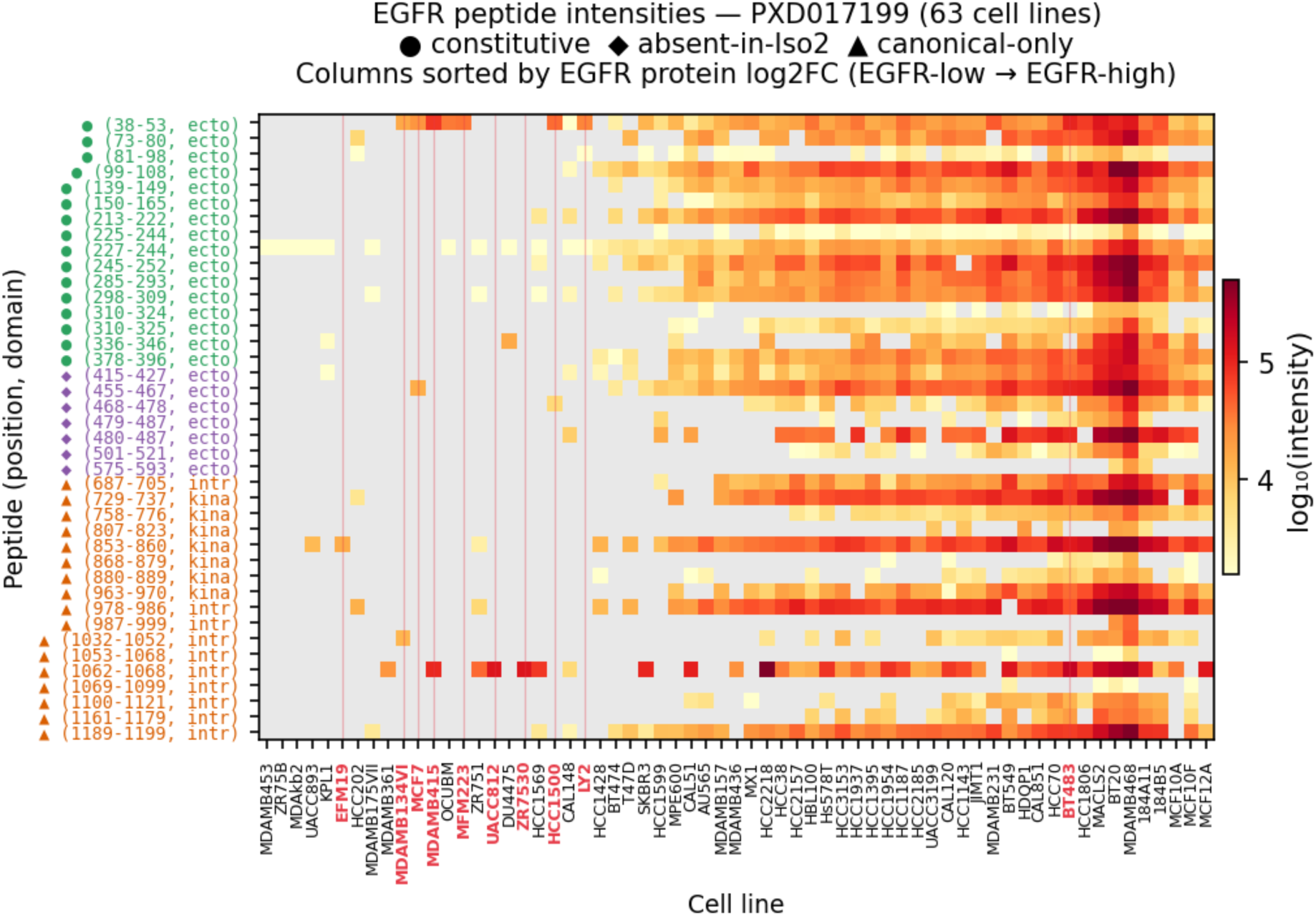
Heatmap showing Spectronaut DIA-MS peptide intensities (log₁₀ PEP.Quantity) for all 40 EGFR (P00533) proteotypic peptides detected in the PXD017199 spectral library across 63 breast cancer cell lines and 5 non-transformed mammary epithelial cell lines. Data were retrieved from ProteomeXchange dataset PXD017199 (Tognetti et al., 2021), which profiled 63 breast cancer cell lines by SWATH-MS using Spectronaut. Rows represent individual tryptic peptides ordered by position along the canonical EGFR sequence (N→C terminus, aa position indicated in row labels). Columns represent cell lines sorted left to right by increasing EGFR protein-level. Grey cells indicate peptides not detected in that cell line. Row symbols denote isoform-class assignment based on mapping to UniProt-annotated alternative isoforms: ● constitutive (aa 38–396; present in canonical and all known alternative isoforms including soluble sEGFR/Iso2); ◆ absent in Iso2 (aa 415–593; present in canonical and membrane-anchored isoforms 3/4, but absent in the truncated soluble ectodomain isoform 2); ▲ canonical-specific (aa 687–1199; intracellular and kinase domain peptides absent from all three alternative isoforms). Cell lines highlighted in red have supposedly high EGFR protein abundance (log_2_FC > –2 relative to normal cell lines) and low EGFR transcript expression (log-FKPM < 2).

**Figure S8.**
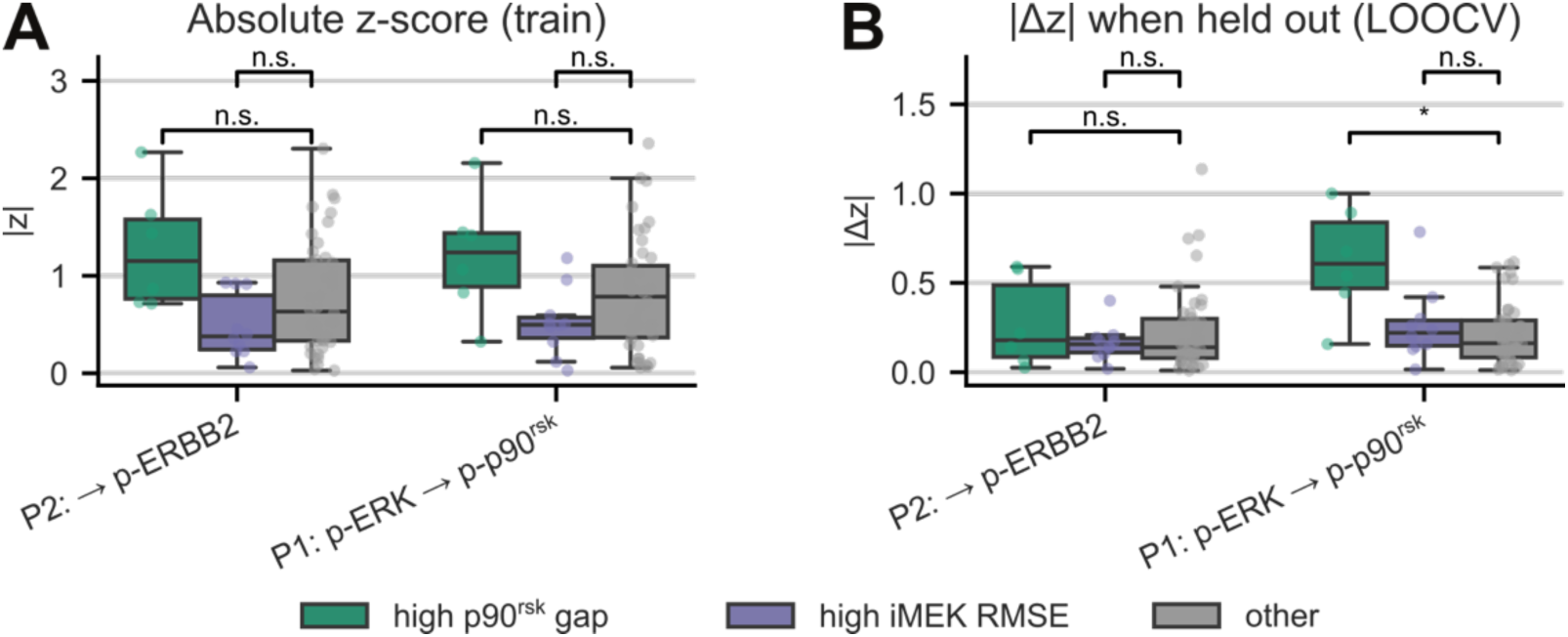
Inferred heterogeneity axes parameters for different high error and gap groups. **(A)** Boxplot of absolute average z-scores of parameter deviations (across n=10 training replicates). **(B)** Boxplot of absolute value of average change in z-scores when cell lines are held out during leave-one-out-cross-validation (LOOCV).

**Figure S9.**
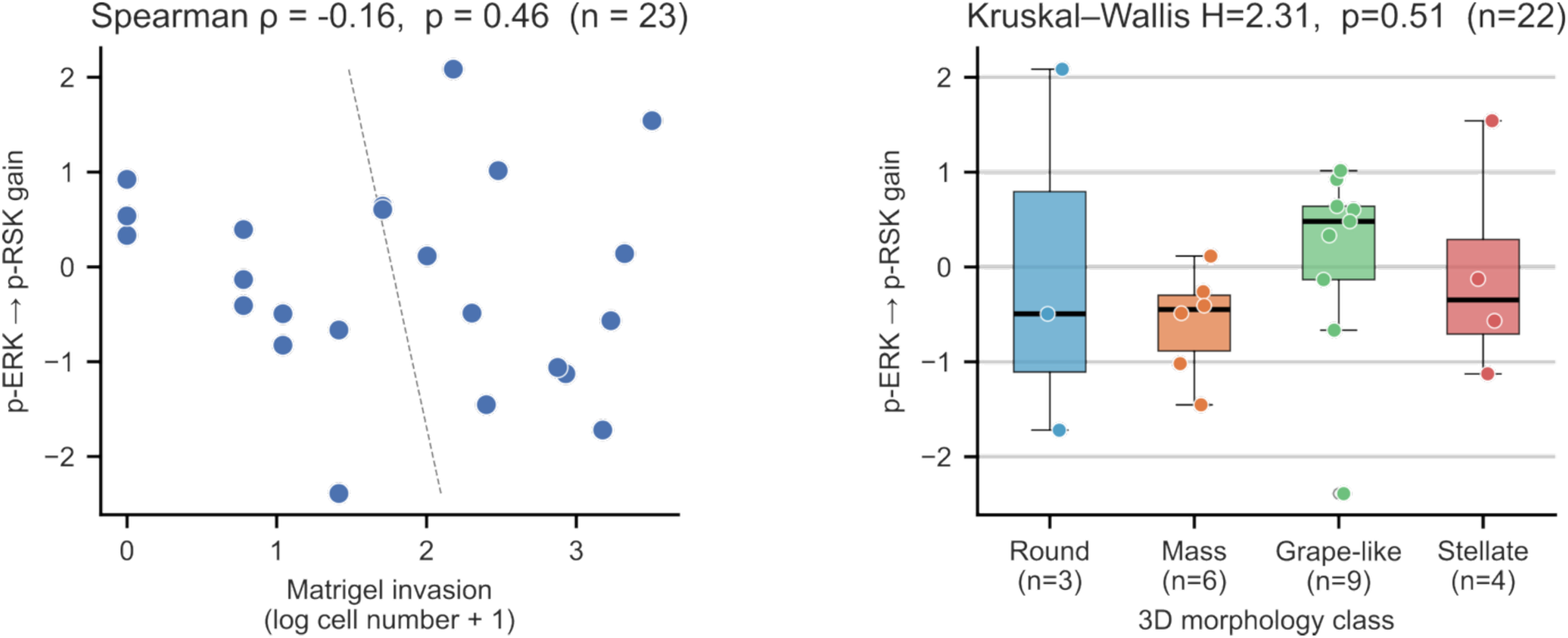
Inferred p-ERK→p-p90^rsk^ gain compared to Matrigel invasion and 3D morphology across breast cancer cell lines. **A**) Scatter plot comparing model-inferred ERK→p90^rsk^ gain parameters (z-scored across cell lines) against Matrigel Boyden chamber invasion (log₁₀ cell number + 1) for breast cancer cell lines with paired data (n = 23). Invasion cell counts (72 h, 8 μm pore Matrigel-coated chambers, 75,000 seeded cells) were extracted from (Neve et al., 2006) (Figure 6B). **(B)** Box plot comparing model-inferred ERK→p90^rsk^ gain parameters (z-scored across cell lines) across different 3D morphology classes as characterised by (Kenny et al., 2007).

